# Genetic context of transgene insertion can influence neurodevelopment in zebrafish

**DOI:** 10.1101/2025.02.28.640904

**Authors:** Anna J. Moyer, Alexia Barcus, Mary E.S. Capps, Jessica A. Chrabasz, Robert L. Lalonde, Christian Mosimann, Summer B. Thyme

## Abstract

The Gal4/*UAS* system is used across model organisms to overexpress target genes in precise cell types and relies on generating transgenic Gal4 driver lines. In zebrafish, the *Tg(elavl3:KalTA4)* (*HuC*) Gal4 line drives robust expression in neurons. We observed an increased prevalence of swim bladder defects in *Tg(elavl3:KalTA4)* zebrafish larvae compared to wildtype siblings, which prompted us to investigate whether transgenic larvae display additional neurobehavioral phenotypes. *Tg(elavl3:KalTA4)* larvae showed alterations in brain activity, brain morphology, and behavior, including increased hindbrain size and reduced activity of the cerebellum. Bulk RNA-seq analysis revealed dysregulation of the transcriptome and suggested an increased ratio of neuronal progenitor cells compared to differentiated neurons. To understand whether these phenotypes derive from Gal4 toxicity or from positional effects related to transgenesis, we used economical low-pass whole genome sequencing to map the *Tol2-*mediated insertion site to chromosome eight. Reduced expression of the neighboring gene *gadd45ga*, a known cell cycle regulator, is consistent with increased proliferation and suggests a role for positional effects. Challenges with creating alternative pan-neuronal lines include the length of the *elavl3* promoter (over 8 kb) and random insertion using traditional transgenesis methods. To facilitate the generation of alternative lines, we cloned five neuronal promoters (*atp6v0cb*, smaller *elavl3, rtn1a, sncb*, and *stmn1b*) ranging from 1.7 kb to 4.3 kb and created *KalTA4* lines using *Tol2* and the phiC31 integrase-based *pIGLET* system. Our study highlights the importance of using appropriate genetic controls and interrogating potential positional effects in new transgenic lines.

**ARTICLE SUMMARY:** The Gal4/*UAS* system is a genetic tool that allows researchers to overexpress specific genes in specific cell types by pairing a Gal4 “driver” line with a *UAS* “reporter” line. In zebrafish, more than 500 transgenic Gal4 lines have been generated using transgenesis methods that insert DNA into random sites in the genome. Here, the authors report that a pan-neuronal Gal4 zebrafish line, *Tg(elavl3:KalTA4)*, has neurobehavioral phenotypes, including changes in brain activity, brain structure, and behavior. These findings emphasize the importance of using appropriate controls and highlight the utility of methods for targeted transgenesis.

## INTRODUCTION

The Gal4/UAS system is a two-part genetic tool originally derived from yeast, where the transcription factor Gal4 binds to *Upstream Activating Sequences* (*UAS*) to drive expression of genes related to galactose metabolism (Giniger et al. 1985). The mechanism for transcriptional activation is conserved across eukaryotes, and model organism researchers have co-opted the Gal4/*UAS* system to drive expression of target genes in specific cell types and developmental windows. Although the simplest experimental design involves placing Gal4 downstream of a cell-type-specific promoter and a target gene downstream of *UAS* sites, the tool has been adapted for diverse zebrafish applications ranging from large-scale enhancer trapping (Asakawa et al. 2008; Marquart et al. 2015; Scott et al. 2007) to interrogation of developmental signaling pathways (Davey et al. 2016; Scheer et al. 2001; Takamiya and Campos-Ortega 2006) to identification of neurons involved in fear conditioning (Lal et al. 2024; Palumbo et al. 2020).

Despite widespread adoption in zebrafish, technical challenges have necessitated modifications to both Gal4 and *UAS* transgenes. Low levels of transcriptional activation can be overcome with a modified version of Gal4 that fuses the DNA-binding domain of Gal4 to the potent viral transcriptional activator VP16 (Davison et al. 2007; Sadowski et al. 1988). However, Gal4 and Gal4-VP16 toxicity, hypothesized to act via squelching of transcriptional machinery, has been reported in yeast (Gill and Ptashne 1988), *Drosophila* (Kramer and Staveley 2003), mouse (Habets et al. 2003), and zebrafish (Akitake et al. 2011; Koster and Fraser 2001). Gal4ff, which contains the DNA-binding domain of Gal4 fused to two fragments of VP16, is less toxic to zebrafish than the original Gal4-VP16 construct (Akitake et al. 2011; Asakawa et al. 2008). An alternative construct, KalTA4, also contains a fragment of VP16 but is further codon optimized for zebrafish. Generating and maintaining *UAS* lines can also present a challenge because silencing of transgenes, especially the short tandem repeats that make up *UAS* constructs and contain methylation-prone CpG sequences, can lead to variegated expression that differs both within and between generations (Akitake et al. 2011; Shoenhard and Granato 2023).

In addition to Gal4 toxicity and silencing of *UAS* repeats, positional effects from random transgene insertion using *Tol2*-mediated transposition can cause variation in expression between lines and mutation of nearby genes or regulatory elements (Nagayoshi et al. 2008). Mutation resulting from random transgenesis is an established issue across model organisms; for example, a mouse line expressing Cre in brown fat possesses major developmental abnormalities (Halurkar et al. 2025). A promising alternative strategy involves integrase–based recombination into known safe harbor sites, which was recently implemented using phiC31 integrase in the *pIGLET* system (Lalonde et al. 2024; Mosimann et al. 2013; Roberts et al. 2014). This approach yields single insertions of transgenes at previously validated genomic loci, avoiding unintended disruption of nearby genes and the labor-intensive outcrossing process required to generate single insertions using *Tol2-*mediated transgenesis. However, zebrafish are highly polymorphic, and even targeted transgenesis may not avoid changes in gene expression caused when the transgene is in linkage disequilibrium with naturally-occurring variants (White et al. 2022).

The *Tg(elavl3:KalTA4)* zebrafish line (*zf562Tg*) was produced with *Tol2* and contains fragments of the *elavl3* promoter, first exon, first intron, and second exon (totaling 8.8 kb) upstream of *KalTA4* (Kim et al. 2014). The strain drives robust expression of target genes in larval neurons and was originally used to evaluate activity of the calcium indicator GCaMP3 during spontaneous eye movements. However, the insertion site was not mapped, *elavl3* expression decreases across the lifetime, and the effects of KalTA4 expression on neurodevelopment have not been assessed. Here, we report multiple neurobehavioral phenotypes in *Tg(elavl3:KalTA4)*^*zf562Tg*^ larvae, map the corresponding insertion site, and uncover changes in expression of the neighboring gene *gadd45ga* that are associated with the *zf562Tg* integration. Inspired by the recent identification of an ultra-short neuronal promoter in mammals (Wang et al. 2023) and to facilitate the generation of pan-neuronal lines that bypass the Gal4/*UAS* system, we also cloned putative neuronal promoters with predicted expression in adult neurons and produced transgenic lines using both *Tol2* and phiC31.

## RESULTS

### *Tg(elavl3:KalTA4)* larvae possess morphological and neurobehavioral phenotypes

We observed a high incidence of morphological defects in the progeny of *Tg(elavl3:KalTA4)* animals and quantified the proportion of larvae lacking inflated swim bladders. At 6 days post-fertilization (dpf), larvae with a single copy of the *elavl3:KalTA4* transgene displayed a significantly higher incidence of swim bladder defects than their wildtype siblings (Fig. 1a–b), while the ratio of transgenic to wildtype larvae did not differ from expected (n=447, P=0.48, chi-square test). To test whether *Tg(elavl3:KalTA4)* larvae show changes in brain activity or structure, we performed phosphorylated Erk (pErk) brain activity mapping, in which pErk and total Erk (tErk) immunostaining is registered to a reference brain atlas (Randlett et al. 2015; Thyme et al. 2019). Comparison of pErk/tERK signal between *Tg(elavl3:KalTA4)* and wildtype siblings, representing changes in brain activity, revealed decreased activity in the cerebellum and increased activity in the rhombencephalon across three independent experiments (Fig. 1c and Supplementary Fig. S1). Deformation during image registration indicates changes in brain structure (Jefferis et al. 2007; Thyme et al. 2019), and compared to their wildtype siblings, *Tg(elavl3:KalTA4)* larvae showed a significant increase in hindbrain size across three independent clutches (Fig. 1c, Supplementary Fig. S1a).

**Fig. 1.**
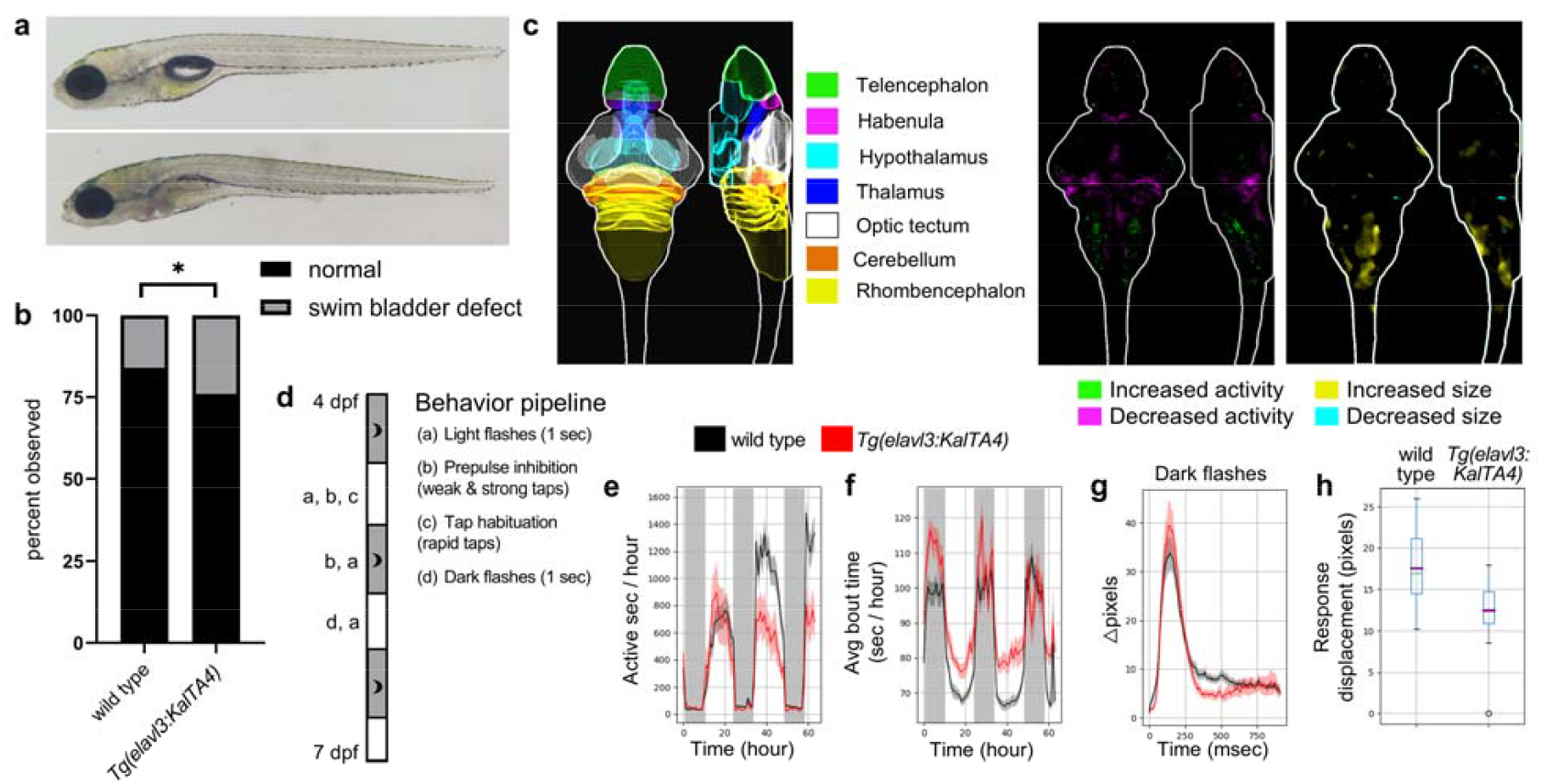
Morphological, brain imaging, and behavioral phenotypes in *Tg(elavl3:KalTA4)* larvae. a) Representative 6 dpf *Tg(elavl3:KalTA4)* transgenic larva lacking inflated swim bladder (bottom) and wildtype sibling with inflated swim bladder (top). b) Quantification of swim bladder defect phenotype in *Tg(elavl3:KalTA4)* transgenic larvae and wildtype siblings. n=447 larvae from five independent clutches. Statistics, chi-square test where *P ≤ 0.05. c) Schematic of seven major brain regions derived from the Z-Brain atlas (left) (Randlett et al. 2015). Brain activity (middle) and structure (right) maps comparing *Tg(elavl3:KalTA4)* transgenic larvae to wildtype siblings. Two additional biological replicates are in Supplementary Fig. S1. n= 18 transgenic and 20 wildtype siblings. d) Schematic of behavioral pipeline from 4 dpf to 7 dpf with stimuli labeled (a) through (d). e) Frequency of motion plot, active seconds binned per hour over the entire experiment (Kruskal-Wallis ANOVA, P=0.004). f) The average time of individual bout motions, binned per hour over the entire experiment (Kruskal-Wallis ANOVA, P=0.021). g) Average movement (pixel-based) of larvae in the mutant and control groups for events where a response to a dark flash was observed. This data is from all dark flashes. h) The response displacement for a dark flash response averaged for each zebrafish from all dark flashes (Kruskal-Wallis ANOVA, P=0.00014). Additional plots of repeatable phenotypes and a second biological replicate are in Supplementary Fig. S1. n= 17 transgenic and 31 wildtype siblings.

We also subjected larvae to a behavioral battery from 4 dpf to 7 dpf, which includes both baseline activity and response to acoustic and visual stimuli (Fig. 1d) (Joo et al. 2020). Both baseline and stimulus-driven behavioral phenotypes were identified (Supplementary Fig. S1b-f). The mutants displayed reduced movement frequency at 6 and 7 dpf (Fig. 1e, Supplementary Fig. S1f), increased bout magnitude as measured by both time of the bout and its displacement (Fig. 1f, Supplementary Fig. S1c, Supplementary Fig. S1d, Supplementary Fig. S1f), and a subtle but consistent change to the dark flash response (Fig. 1g, Fig. 1h, Supplementary Fig. S1e, Supplementary Fig. S1f).

### Transcriptional profiling reveals abnormal proliferation and cell type composition in *Tg(elavl3:KalTA4)* larvae

To probe the molecular underpinnings of brain imaging and behavioral phenotypes, we performed bulk RNA-seq of 6 dpf heads from *Tg(elavl3:KalTA4)* animals and wildtype siblings (n=4 transgenic and 4 wild type). Compared to control, *Tg(elavl3:KalTA4)* samples showed significant dysregulation of the transcriptome, with downregulation of 111 genes and upregulation of 223 at a threshold of P adjusted (padj) <0.05 (Fig. 2a and Supplementary Table 1). Gene ontology (GO) analysis of up and down-regulated genes identified terms related to cell cycle and growth hormone receptor signaling among upregulated genes and terms related to synapse organization among downregulated genes (Supplementary Fig. S2 and Supplementary Tables 2-3). Similarly, Gene Set Enrichment Analysis (GSEA) using the C5 Molecular Signatures Database revealed positive enrichment of terms related to cell cycle, mitotic spindle, DNA replication, chromatin organization, and splicing, and negative enrichment of terms related to synapses and ions/channels (Fig. 2b and Supplementary Table 4). Genes contributing to enrichment of these terms included canonical cell cycle regulators such as *ccna2, aurkb*, and *bub1ba* (Fig. 2c). GSEA using gene sets derived from single-cell RNA-seq data can suggest changes in cell state, and GSEA with terms generated using previously published single-cell data from 5 dpf heads (Raj et al. 2020) revealed positive enrichment of progenitors and negative enrichment of mature neuronal subtypes (Fig. 2d and Supplementary Table 5). Consistent with bulk RNA-seq data, 6 dpf *Tg(elavl3:KalTA4)* larvae showed an increased number of phospho-histone H3 (pHH3) foci in the brain and spinal cord (Fig. 2e-f). Together, these analyses indicate increased proliferation and reduced expression of genes related to differentiation and synaptic development.

**Fig. 2.**
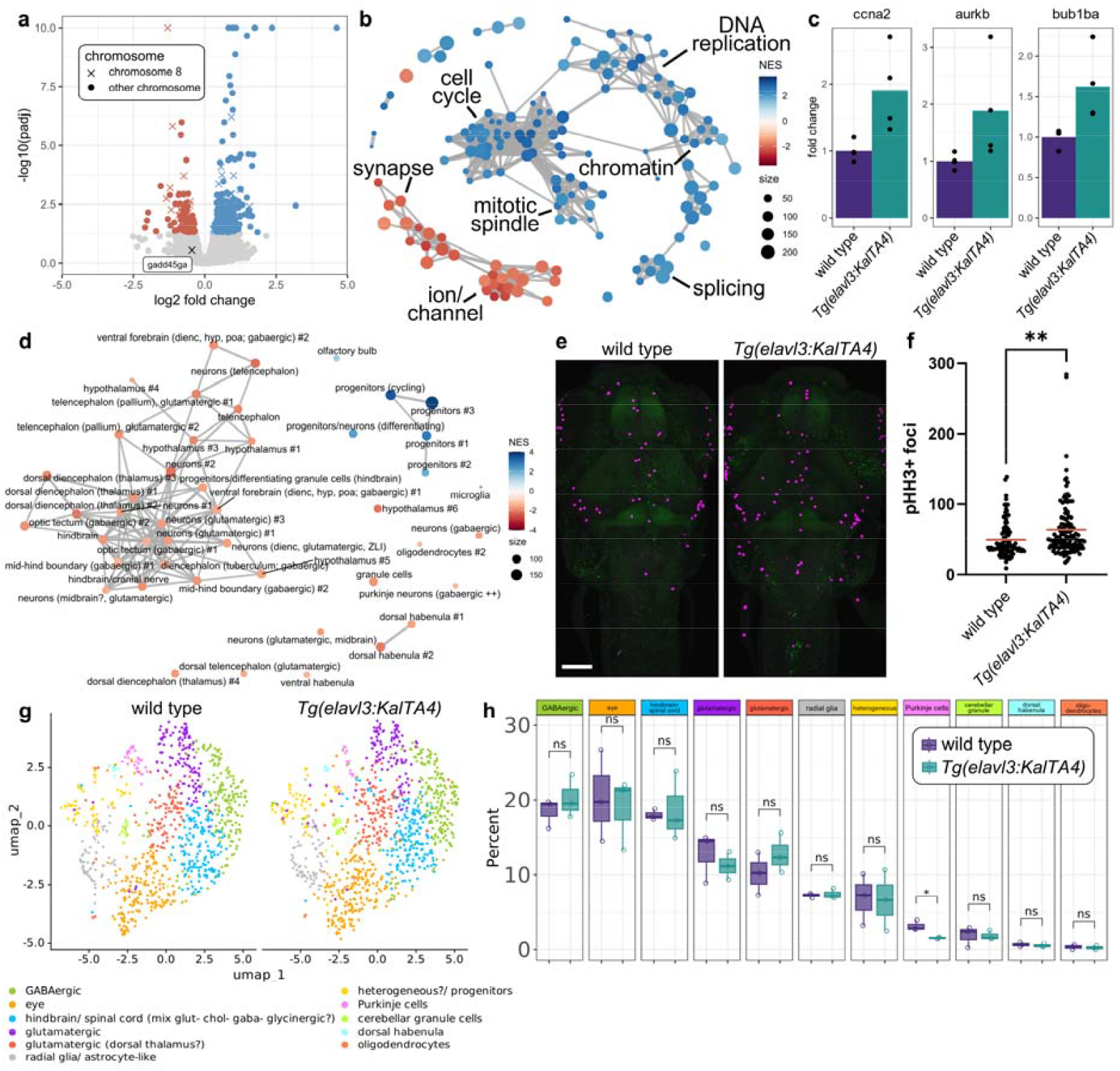
Bulk and sci-RNA-seq reveal changes in proliferation and cell-type composition. a) Volcano plot of 6 dpf RNA-seq, with upregulated genes labeled in blue and downregulated genes labeled in red (padj <0.05). Y-axis values over ten are plotted at ten. X denotes transcripts encoded by chromosome eight genes and circles denote all other chromosomes. b) Network plot with top 150 terms identified by C5 GSEA. Nodes represent GSEA terms and edges represent genes shared between terms. Node size represents term size and color represents the normalized enrichment score. c) Bar plots of normalized RNA-seq counts for selected genes. P values for all genes are in Supplementary Table 1. d) Network plot with terms identified by GSEA using terms generated from previously published single-cell data (Raj et al. 2020). Nodes represent GSEA terms and edges represent genes shared between terms. Node size represents term size and color represents the normalized enrichment score. e) Representative maximum intensity projections of confocal images for 6 dpf larvae labeled with pHH3 (magenta) and tERK (green). Images shown are 145-slice maximal projections with 3 μm interval. Scale bar = 100 μm. f) Quantification of pHH3+ foci. Red line represents average. n=93 wild type and 119 *Tg(elavl3:KalTa4)* animals from four independent clutches. g) Uniform Manifold Approximation and Projection (UMAP) representation of eleven neuronal/glial clusters from wildtype nuclei (left) and *Tg(elavl3:KalTA4)* nuclei (right). h) Box plot representing the percentage of cells in each neuronal/glial cluster in wild type and *Tg(elavl3:KalTa4)*. n=3 wild type and 3 *Tg(elavl3:KalTa4)*. Statistics, two-tailed t-test where *P ≤ 0.05; **P ≤ 0.01.

We next used single cell combinatorial indexing RNA sequencing (sci-RNA-seq) to explore changes to the transcriptome at the single-nucleus level (Martin et al. 2023). Initial clustering of 1452 nuclei from wild type 6 dpf heads and 1517 nuclei from *Tg(elavl3:KalTA4)* heads defined ten non-neuronal/eye clusters and six neuronal/glial clusters (Supplementary Fig. S3 and Supplementary Table 6), which we sub-clustered to identify eleven neuronal/glial subtypes (Fig. 2g, Supplementary Fig. S4, and Supplementary Table 7). Although *Tg(elavl3:KalTA4)* heads were not missing any major cell types compared to wildtype samples, we observed a small decrease in the proportion of cerebellar Purkinje cells (Fig. 2h).

### Larvae with two copies of *elavl3:KalTa4* insertion have more severe phenotypes than larvae with one insertion

Having established that *Tg(elavl3:KalTA4)* larvae possess brain imaging, behavior, and transcriptional phenotypes, we sought to understand whether these phenotypes derive from insertional mutagenesis or from overexpression of *KalTA4*. Naturally-occurring variants that are in linkage disequilibrium with mutations can cause changes in transcription that manifest in bulk RNA-seq data as enrichment of differentially expressed genes across a single chromosome (Capps et al. 2024; White et al. 2022). We observed a significant enrichment of differentially expressed genes on chromosome eight (Fig. 3a and Fig. 2a), and low-coverage whole genome sequencing confirmed that the *elavl3:KalTA4* transgene maps to an intergenic region of chromosome eight between *unm_hu7912* and *gadd45ga* (Fig. 3b). Mapping enabled unambiguous genotyping of larvae with two copies of the *KalTA4* transgene, and larvae with two copies possessed significantly more swim bladder defects (Fig. 3c) and brain imaging phenotypes (Fig. 3d) than larvae with one copy. Bulk RNA-seq data indicated possible downregulation of the neighboring gene *gadd45ga* (Supplementary Table 1), the mammalian orthologs of which regulate the cell cycle via inhibition of Cdk1/cyclinB1 (Vairapandi et al. 2002). qPCR confirmed a significant reduction in *gadd45ga* expression, with 71% and 58% of wild type in larvae with one or two copies of the *elavl3:KalTA4* transgene, respectively (Fig. 3e).

**Fig. 3.**
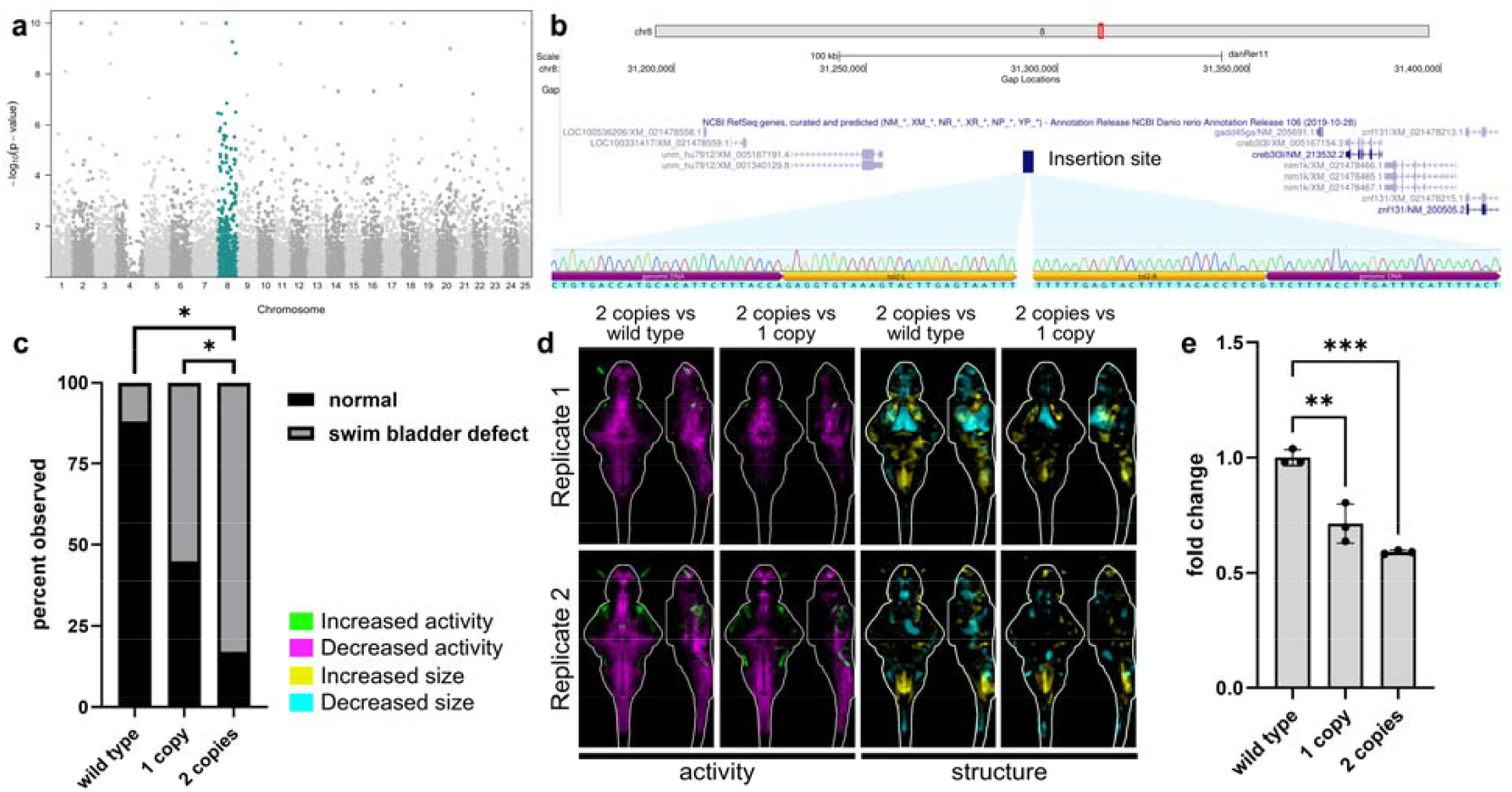
*elavl3:KalTA4* transgene maps to zebrafish chromosome eight. a) Manhattan plot of bulk RNA-seq P-values plotted in chromosomal order. Transcripts encoded by chromosome eight are significantly enriched using GSEA (teal). b) Schematic of insertion location with NCBI RefSeq genes visualized using the UCSC Genome Browser (GRCz11/danRer11). Sanger sequencing shown spanning genomic DNA and *Tol2* arms. c) Quantification of swim bladder defect phenotype in larvae with one or two copies of *elavl3:KalTA4* and wildtype siblings. n=83 larvae from three independent clutches. Statistics, pairwise chi-square test where *P ≤ 0.05; ****P≤ 0.0001. d) Brain activity and structure maps comparing larvae with two copies of the *elavl3:KalTA4* transgene to wildtype siblings and to larvae with one copy. Images show sum-of-slices intensity projection inside the brain (white outline) for two replicates. n= 12 with two copies, 24 with one copy, and 22 wildtype siblings for replicate one and n= 20 with two copies, 37 with one copy, and 17 wildtype siblings for replicate two. e) qPCR showing relative fold change of *gadd45ga* in 6 dpf heads. n=3 per genotype. Statistics, one-way ANOVA with Dunnett’s multiple comparisons test. **P ≤ 0.01; ***P≤ 0.001.

### Generation of alternative Gal4 lines with pan-neuronal expression

To understand whether expression of *KalTA4* in neurons is sufficient to cause neurodevelopmental phenotypes, we next sought to generate *KalTA4* lines with comparably short promoters driving pan-neuronal expression across the lifetime. We selected four genes with high correlation (r) to *elavl3* expression in the Daniocell single cell atlas, which surveys gene expression in whole animals up to 5 dpf (Fig. 4a) (Sur et al. 2023). Like *elavl3*, expression of *stmn1b, rtn1b, sncb* and *atp6v0cb* appears pan-neuronal in Daniocell (Fig. 4b). To enrich for promoters that maintain expression throughout the lifetime, we also examined previously published forebrain single-cell data from 6 dpf, 15 dpf, and adult forebrain (Pandey et al. 2023). Expression of *elavl3* peaks in committed precursors and shows lower expression in adult neurons, whereas expression of *stmn1b, rtn1b, sncb*, and *atp6v0cb* appears to be maintained in the adult brain (Supplementary Fig. S5).

**Fig. 4.**
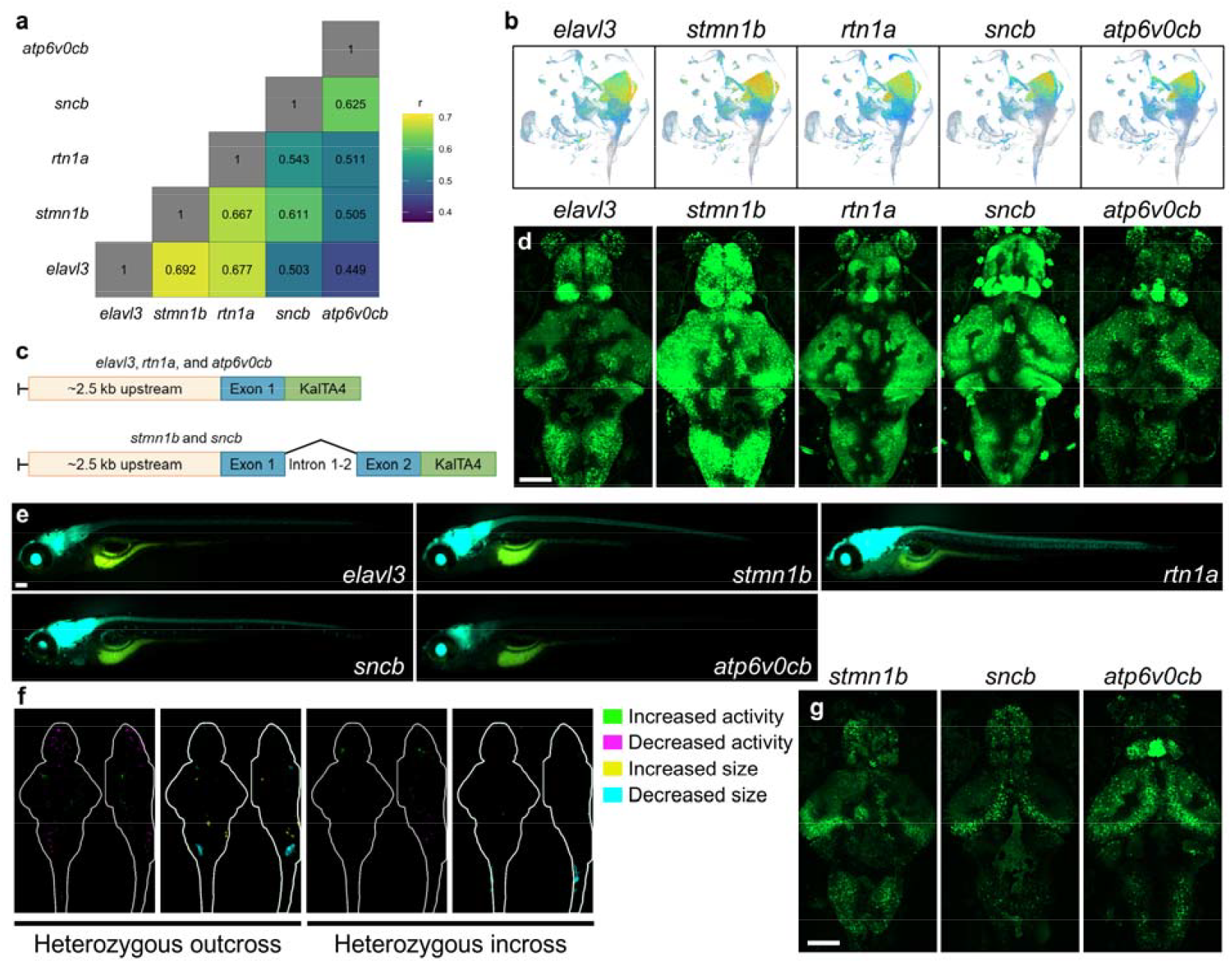
Generation of transgenic lines with pan-neuronal expression of *KalTA4* a) Correlation matrix showing pairwise Spearman correlation (r) between expression of five genes in the Daniocell single cell atlas (https://daniocell.nichd.nih.gov/), which comprises whole animals from 3 hours post-fertilization to 5 dpf (Sur et al. 2023). b) Uniform Manifold Approximation and Projection (UMAP) representation of expression of *elavl3, stmn1b, rtn1a, sncb*, and *atp6v0cb* obtained from the Daniocell single cell atlas. Detectable expression is represented by a gradient from blue (low) to red (high). c) Schematic of promoter constructs cloned into dual *Tol2*/phiC31-compatible plasmids for *elavl3, rtn1a*, and *atp6v0cb* (top), as well as *stmn1b* and *sncb* (bottom). d) Confocal images of brains from live 6 dpf transgenic larvae generated using *Tol2*. Images shown are 145-slice maximal projections with 3 μm interval. Scale bar = 100 μm. Uncropped images and additional independent lines are shown in Supplementary Fig. S6. e) Live fluorescent images of 6 dpf larvae generated using *Tol2*. Additional independent lines are shown in Supplementary Fig. S2. Scale bar = 100 μm. f) Brain activity and structure maps comparing heterozygous *pIGLET14a* transgenic larvae to wildtype siblings from heterozygous outcross (left, n=29 heterozygous and 27 wild type) and homozygous larvae to heterozygous and wildtype siblings from heterozygous incross (right, n=10 homozygous and 34 heterozygous/wild type). g) Confocal images of brains from live 6 dpf transgenic larvae generated using phiC31. Scale bar = 100 μm.

Although expression of ∼2.8 kb of the *elavl3* promoter is sufficient to drive GFP expression in neurons (Park et al. 2000), the most commonly used promoter construct contains 2.8 kb of the promoter plus the first exon, intron 1-2, and a fragment of exon two, resulting in a ∼9 kb construct (Higashijima et al. 2003). To minimize challenges in cloning and transgenesis efficiency, we tested whether shorter promoter constructs were capable of driving *KalTA4* expression in neurons. For *elavl3, rtn1a*, and *atp6v0cb*, the construct contains the upstream promoter and a fragment of the first exon, with the start codon of *KalTA4* replacing the start codon of the native transcript to create constructs of 2.9 kb, 1.7 kb, and 3 kb, respectively (Fig. 4c). The native start codons of *stmn1b* and *sncb* are located in the second exon, and constructs containing exon one, intron 1-2, and a fragment of exon two were 4.3 kb and 3 kb. All five promoters were capable of driving the expression of *UAS:GFP* in the brain and spinal cord in *Tol2*-generated lines, but we observed significant variation in fluorescence intensity between independent lines (Fig. 4d and e, Supplementary Fig. S6-7). Heterozygous animals with one copy of the empty phiC31 *pIGLET14a* landing site, derived from a heterozygous outcross, and homozygous animals with two copies of the landing site, derived from a heterozygous incross, did not show significant brain activity or structural phenotypes (Fig. 4f, Supplementary Fig. S8), and we used phiC31 to generate *pIGLET14a KalTA4-*driver lines for *stmn1b, sncb*, and *atp6v0cb* (Fig. 4g). In turn, behavioral analysis of a subset of the newly generated *Tol2* and phiC31-generated *KalTA4* lines had normal baseline behavior, indicating that pan-neuronal expression of *KalTA4* is not sufficient to drive changes in baseline larval behavior (Supplementary Fig. S9).

## DISCUSSION

The potential for genomic disruption following random transgenesis methods is shared across model and non-model organisms, from *Tol2* transposition in zebrafish to insertion of bacterial artificial chromosomes in mice. For example, Halurkar et al. recently reported that the commonly used mouse line *Ucp1-Cre* has major phenotypic abnormalities related to the transgene insertion site, which raises concerns about the interpretation of brown fat research conducted using this line (Halurkar et al. 2025). Outbred, highly polymorphic species like zebrafish require additional attention as mutations and transgene insertion sites are in linkage disequilibrium with naturally-occurring variants that affect gene expression (White et al. 2022). The advent of extremely sensitive phenotyping methods, including RNA-seq and brain activity mapping, makes it possible to detect phenotypes caused by allele-specific gene expression rather than by the mutation or insertion of interest, and these effects are compounded in incrossed homozygous animals (Supplementary Fig. S8).

In this study, we characterize the morphological and neurobehavioral phenotypes of *Tg(elavl3:KalTA4)* larvae (Kim et al. 2014). Previous reports have raised concerns about Gal4 toxicity (Akitake et al. 2011; Gill and Ptashne 1988; Habets et al. 2003), and we hypothesized that the phenotypes in *Tg(elavl3:KalTA4)* larvae arise from overexpression of Gal4 or from positional effects related to the insertion site of the transgene. In bulk RNA-seq data, we observed a significant enrichment of differentially expressed genes on chromosome eight, which is consistent with allele-specific expression of genes on the same chromosome as the transgene and suggests positional effects (White et al. 2022) (Fig. 3a). However, some changes in gene expression could also result from Gal4 toxicity or from off-target binding of Gal4 to zebrafish promoters. Further support for a positional effect-mediated impairment comes from qPCR demonstrating the downregulation of the neighboring gene *gadd45ga* (Fig. 3e). We hypothesize that the original *Tol2-*generated insertion disrupts a regulatory element required for normal expression of *gadd45ga* or that the transgene is in linkage disequilibrium with a variant that affects *gadd45ga* expression. Downregulation of mammalian *GADD45g* promotes proliferation in multiple cellular contexts and tumor types (Guo et al. 2021; Ying et al. 2005; Zhang et al. 2014; Zhang et al. 2024). Regardless of the mechanism by which the *KalTA4* insertion affects *gadd45ga* expression, disruption of this transcript is consistent with the increased number of pHH3+ foci observed in *Tg(elavl3:KalTA4)* brains. Downregulation of *gadd45ga* in *Tg(elavl3:KalTA4)* larvae and the lack of baseline behavioral changes in newly generated pan-neuronal Gal4 lines support a model where positional effects, and not Gal4 toxicity, are responsible for the neurobehavioral phenotypes in the original *Tg(elavl3:KalTA4)* line.

Our observations emphasize the importance of validating new zebrafish lines and including appropriate control siblings. When using the Gal4/*UAS* system, the phenotyping of wild type, Gal4-only, and *UAS*-only control siblings should identify potential issues with Gal4 toxicity or genomic disruption from transgene insertion. Validation of shorter pan-neuronal promoters may also facilitate the generation of lines that bypass the potential toxicity of the Gal4/*UAS* system by cloning these promoters directly upstream of candidate ORFs of interest. We identified five putative promoter sequences capable of driving GFP expression in the brain and spinal cord, but the strength of expression varied both between independent transgenic lines and between promoters, with the *atp6v0cb* promoter showing weak expression across all lines. As previously reported, we observed muscle expression in some *Tol2-*generated *elavl3* lines, suggesting that the upstream MyT1 binding site (not included in our construct) is involved in restricting expression to neurons (Park et al. 2000).

Our study also highlights the utility of safe harbor landing sites for transgenesis in zebrafish and other outbred model organisms. Establishing a lack of phenotypes in zebrafish with empty landing sites ensures that phenotypes observed following transgenesis are due to transgene insertion rather than to positional effects; we observed minimal brain structure and activity phenotypes in larvae with one copy of the phiC31-based *pIGLET14a* landing site (Lalonde et al. 2024). Safe harbor sites also offer a possible solution for the allele-specific gene expression observed in RNA-seq data. Comparing two complementary transgenic lines created using zebrafish strains with landing sites on two different chromosomes, such as *pIGLET14a* and *pIGLET24b*, can nominate differentially expressed genes shared between lines. However, like mutant lines, incrossing transgenic zebrafish to produce homozygous animals may still produce phenotypes related to the homozygosity of alleles linked to the landing site loci by their physical proximity (White et al. 2022). Detailed haplotyping of landing site strains is therefore a desirable next step in validating such new reagents.

## MATERIALS AND METHODS

Additional methods are available in Supplementary Materials and Methods. Animal experiments were approved by the UMass Chan Institutional Animal Care and Use Committee (IACUC protocol 202300000053) and UAB Institutional Animal Care and Use Committee (IACUC protocols 22155/21744). Phosphorylated-ERK (pErk)/total Erk (tERK) immunostaining was performed as previously described (Capps et al. 2024). Behavioral assays were performed using custom-built behavioral rigs as previously described (Joo et al. 2020; Thyme et al. 2019). The bulk RNA-seq protocol was performed as previously described (Capps et al. 2024) and is based on the SMART-Seq2 protocol (Trombetta et al. 2014). Sci-RNA-seq was performed following the “Tiny-Sci” protocol (Martin et al. 2023). Analysis pipeline for bulk RNA-seq is available on GitHub (https://github.com/thymelab/BulkRNASeq) and in the Supplementary Materials and Methods, and analysis pipeline for Sci-RNA-seq was performed according to Martin et al. (https://github.com/bethmartin/sci-RNA-seq3_pipeline/tree/master) (2023). The *elavl3:KalTA4* transgene insertion was mapped using low-pass whole genome sequencing of genomic DNA (15 Gb, ∼10x) by Azenta Life Sciences. Quantitative reverse transcription PCR (qPCR) was performed as previously described (Capps et al. 2024) with the *rpl13a* gene as the reference (Rassier et al. 2020). Putative pan-neuronal promoters were purified from genomic DNA and cloned into dual-*Tol2*/phiC31-compatible backbone using Gibson assembly. Plasmid maps are available from Zenodo.

## DATA AVAILABILITY

Plasmids are available upon request and will be deposited to Addgene. Bulk RNA-seq (GEO GSE288241), low-pass whole genome sequencing (SRA SRP560737), and sci-RNA-seq (GEO GSE288775) data have been deposited in publicly available databases. Additional code and data are available on Zenodo: 10.5281/zenodo.14751662

## ACKNOWLEDGEMENTS

The authors thank Dr. Michael Crowley at the UAB Heflin Center for Genomic Science Core Laboratories, UMass Chan and UAB zebrafish facility staff, and the UMass Chan High Performance Computing cluster for supporting this research. We also thank Dr. Nathan Lawson for providing the *pcDNA3*.*1 phiC31* plasmid, Dr. Koichi Kawakami for providing *Tg(5xUAS:EGFP)* zebrafish, and research technicians Claire Conklin, Camden Cummings, Morgan Klein, and Graham Latino for experimental support.

## LITERATURE CITED

Akitake CM, Macurak M, Halpern ME, Goll MG. 2011. Transgenerational analysis of transcriptional silencing in zebrafish. Dev Biol. 352(2):191–201.

Asakawa K, Suster ML, Mizusawa K, Nagayoshi S, Kotani T, Urasaki A, Kishimoto Y, Hibi M, Kawakami K. 2008. Genetic dissection of neural circuits by tol2 transposon-mediated gal4 gene and enhancer trapping in zebrafish. Proc Natl Acad Sci U S A. 105(4):1255–1260.

Bischof J, Maeda RK, Hediger M, Karch F, Basler K. 2007. An optimized transgenesis system for drosophila using germ-line-specific phic31 integrases. Proc Natl Acad Sci U S A. 104(9):3312–3317.

Bolte S, Cordelieres FP. 2006. A guided tour into subcellular colocalization analysis in light microscopy. J Microsc. 224(Pt 3):213–232.

Capps MES, Moyer AJ, Conklin CL, Martina V, Torija-Olson EG, Klein MC, Gannaway WC, Calhoun CCS, Vivian MD, Thyme SB. 2024. Diencephalic and neuropeptidergic dysfunction in zebrafish with autism risk mutations. bioRxiv.2024.2001.2018.576309.

Davey CF, Mathewson AW, Moens CB. 2016. Pcp signaling between migrating neurons and their planar-polarized neuroepithelial environment controls filopodial dynamics and directional migration. PLoS Genet. 12(3):e1005934.

Davison JM, Akitake CM, Goll MG, Rhee JM, Gosse N, Baier H, Halpern ME, Leach SD, Parsons MJ. 2007. Transactivation from gal4-vp16 transgenic insertions for tissue-specific cell labeling and ablation in zebrafish. Dev Biol. 304(2):811–824.

Dobin A, Davis CA, Schlesinger F, Drenkow J, Zaleski C, Jha S, Batut P, Chaisson M, Gingeras TR. 2013. Star: Ultrafast universal rna-seq aligner. Bioinformatics. 29(1):15–21.

Gill G, Ptashne M. 1988. Negative effect of the transcriptional activator gal4. Nature. 334(6184):721–724.

Giniger E, Varnum SM, Ptashne M. 1985. Specific DNA binding of gal4, a positive regulatory protein of yeast. Cell. 40(4):767–774.

Guo D, Zhao Y, Wang N, You N, Zhu W, Zhang P, Ren Q, Yin J, Cheng T, Ma X. 2021. Gadd45g acts as a novel tumor suppressor, and its activation suggests new combination regimens for the treatment of aml. Blood. 138(6):464–479.

Habets PE, Clout DE, Lekanne Deprez RH, van Roon MA, Moorman AF, Christoffels VM. 2003. Cardiac expression of gal4 causes cardiomyopathy in a dose-dependent manner. J Muscle Res Cell Motil. 24(2-3):205–209.

Halurkar MS, Inoue O, Singh A, Mukherjee R, Ginugu M, Ahn C, Bonatto Paese CL, Duszynski M, Brugmann SA, Lim HW et al. 2025. The widely used ucp1-cre transgene elicits complex developmental and metabolic phenotypes. Nat Commun. 16(1):770.

Hao Y, Stuart T, Kowalski MH, Choudhary S, Hoffman P, Hartman A, Srivastava A, Molla G, Madad S, Fernandez-Granda C et al. 2024. Dictionary learning for integrative, multimodal and scalable single-cell analysis. Nat Biotechnol. 42(2):293–304.

Higashijima S, Masino MA, Mandel G, Fetcho JR. 2003. Imaging neuronal activity during zebrafish behavior with a genetically encoded calcium indicator. J Neurophysiol. 90(6):3986–3997.

Jefferis GS, Potter CJ, Chan AM, Marin EC, Rohlfing T, Maurer CR, Jr., Luo L. 2007. Comprehensive maps of drosophila higher olfactory centers: Spatially segregated fruit and pheromone representation. Cell. 128(6):1187–1203.

Joo W, Vivian MD, Graham BJ, Soucy ER, Thyme SB. 2020. A customizable low-cost system for massively parallel zebrafish behavioral phenotyping. Front Behav Neurosci. 14:606900.

Kawakami K, Takeda H, Kawakami N, Kobayashi M, Matsuda N, Mishina M. 2004. A transposon-mediated gene trap approach identifies developmentally regulated genes in zebrafish. Dev Cell. 7(1):133–144.

Kim CK, Miri A, Leung LC, Berndt A, Mourrain P, Tank DW, Burdine RD. 2014. Prolonged, brain-wide expression of nuclear-localized gcamp3 for functional circuit mapping. Front Neural Circuits. 8:138.

Kolberg L, Raudvere U, Kuzmin I, Vilo J, Peterson H. 2020. Gprofiler2 --an r package for gene list functional enrichment analysis and namespace conversion toolset g:Profiler. F1000Res. 9.

Koster RW, Fraser SE. 2001. Tracing transgene expression in living zebrafish embryos. Dev Biol. 233(2):329–346.

Kramer JM, Staveley BE. 2003. Gal4 causes developmental defects and apoptosis when expressed in the developing eye of drosophila melanogaster. Genet Mol Res. 2(1):43–47.

Lal P, Tanabe H, Kawakami K. 2024. Genetic identification of neural circuits essential for active avoidance fear conditioning in adult zebrafish. Methods Mol Biol. 2707:169–181.

Lalonde RL, Wells HH, Kemmler CL, Nieuwenhuize S, Lerma R, Burger A, Mosimann C. 2024. Piglet: Safe harbor landing sites for reproducible and efficient transgenesis in zebrafish. Sci Adv. 10(23):eadn6603.

Lawson ND, Li R, Shin M, Grosse A, Yukselen O, Stone OA, Kucukural A, Zhu L. 2020. An improved zebrafish transcriptome annotation for sensitive and comprehensive detection of cell type-specific genes. Elife. 9.

Liberzon A, Birger C, Thorvaldsdottir H, Ghandi M, Mesirov JP, Tamayo P. 2015. The molecular signatures database (msigdb) hallmark gene set collection. Cell Syst. 1(6):417–425.

Love MI, Huber W, Anders S. 2014. Moderated estimation of fold change and dispersion for rna-seq data with deseq2. Genome Biol. 15(12):550.

Marquart GD, Tabor KM, Brown M, Strykowski JL, Varshney GK, LaFave MC, Mueller T, Burgess SM, Higashijima S, Burgess HA. 2015. A 3d searchable database of transgenic zebrafish gal4 and cre lines for functional neuroanatomy studies. Front Neural Circuits. 9:78.

Martin BK, Qiu C, Nichols E, Phung M, Green-Gladden R, Srivatsan S, Blecher-Gonen R, Beliveau BJ, Trapnell C, Cao J et al. 2023. Optimized single-nucleus transcriptional profiling by combinatorial indexing. Nat Protoc. 18(1):188–207.

Mosimann C, Puller AC, Lawson KL, Tschopp P, Amsterdam A, Zon LI. 2013. Site-directed zebrafish transgenesis into single landing sites with the phic31 integrase system. Dev Dyn. 242(8):949–963.

Nagayoshi S, Hayashi E, Abe G, Osato N, Asakawa K, Urasaki A, Horikawa K, Ikeo K, Takeda H, Kawakami K. 2008. Insertional mutagenesis by the tol2 transposon-mediated enhancer trap approach generated mutations in two developmental genes: Tcf7 and synembryn-like. Development. 135(1):159–169.

Palumbo F, Serneels B, Pelgrims R, Yaksi E. 2020. The zebrafish dorsolateral habenula is required for updating learned behaviors. Cell Rep. 32(8):108054.

Pandey S, Moyer AJ, Thyme SB. 2023. A single-cell transcriptome atlas of the maturing zebrafish telencephalon. Genome Res. 33(4):658–671.

Pandey S, Shekhar K, Regev A, Schier AF. 2018. Comprehensive identification and spatial mapping of habenular neuronal types using single-cell rna-seq. Curr Biol. 28(7):1052–1065 e1057.

Park HC, Kim CH, Bae YK, Yeo SY, Kim SH, Hong SK, Shin J, Yoo KW, Hibi M, Hirano T et al. 2000. Analysis of upstream elements in the huc promoter leads to the establishment of transgenic zebrafish with fluorescent neurons. Dev Biol. 227(2):279–293.

Raj B, Farrell JA, Liu J, El Kholtei J, Carte AN, Navajas Acedo J, Du LY, McKenna A, Relic D, Leslie JM et al. 2020. Emergence of neuronal diversity during vertebrate brain development. Neuron. 108(6):1058–1074 e1056.

Randlett O, Wee CL, Naumann EA, Nnaemeka O, Schoppik D, Fitzgerald JE, Portugues R, Lacoste AM, Riegler C, Engert F et al. 2015. Whole-brain activity mapping onto a zebrafish brain atlas. Nat Methods. 12(11):1039–1046.

Rassier GT, Silveira TLR, Remiao MH, Daneluz LO, Martins AWS, Dellagostin EN, Ortiz HG, Domingues WB, Komninou ER, Kutter MT et al. 2020. Evaluation of qpcr reference genes in gh-overexpressing transgenic zebrafish (danio rerio). Sci Rep. 10(1):12692.

Roberts JA, Miguel-Escalada I, Slovik KJ, Walsh KT, Hadzhiev Y, Sanges R, Stupka E, Marsh EK, Balciuniene J, Balciunas D et al. 2014. Targeted transgene integration overcomes variability of position effects in zebrafish. Development. 141(3):715–724.

Sadowski I, Ma J, Triezenberg S, Ptashne M. 1988. Gal4-vp16 is an unusually potent transcriptional activator. Nature. 335(6190):563–564.

Scheer N, Groth A, Hans S, Campos-Ortega JA. 2001. An instructive function for notch in promoting gliogenesis in the zebrafish retina. Development. 128(7):1099–1107.

Scott EK, Mason L, Arrenberg AB, Ziv L, Gosse NJ, Xiao T, Chi NC, Asakawa K, Kawakami K, Baier H. 2007. Targeting neural circuitry in zebrafish using gal4 enhancer trapping. Nat Methods. 4(4):323–326.

Shoenhard H, Granato M. 2023. Multivariate analysis of variegated expression in neurons: A strategy for unbiased localization of gene function to candidate brain regions in larval zebrafish. PLoS One. 18(2):e0281609.

Sur A, Wang Y, Capar P, Margolin G, Prochaska MK, Farrell JA. 2023. Single-cell analysis of shared signatures and transcriptional diversity during zebrafish development. Dev Cell. 58(24):3028–3047 e3012.

Takamiya M, Campos-Ortega JA. 2006. Hedgehog signalling controls zebrafish neural keel morphogenesis via its level-dependent effects on neurogenesis. Dev Dyn. 235(4):978–997.

Thyme SB, Pieper LM, Li EH, Pandey S, Wang Y, Morris NS, Sha C, Choi JW, Herrera KJ, Soucy ER et al. 2019. Phenotypic landscape of schizophrenia-associated genes defines candidates and their shared functions. Cell. 177(2):478–491 e420.

Trombetta JJ, Gennert D, Lu D, Satija R, Shalek AK, Regev A. 2014. Preparation of single-cell rna-seq libraries for next generation sequencing. Curr Protoc Mol Biol. 107:4 22 21–24 22 17.

Vairapandi M, Balliet AG, Hoffman B, Liebermann DA. 2002. Gadd45b and gadd45g are cdc2/cyclinb1 kinase inhibitors with a role in s and g2/m cell cycle checkpoints induced by genotoxic stress. J Cell Physiol. 192(3):327–338.

Wang J, Lin J, Chen Y, Liu J, Zheng Q, Deng M, Wang R, Zhang Y, Feng S, Xu Z et al. 2023. An ultra-compact promoter drives widespread neuronal expression in mouse and monkey brains. Cell Rep. 42(11):113348.

White RJ, Mackay E, Wilson SW, Busch-Nentwich EM. 2022. Allele-specific gene expression can underlie altered transcript abundance in zebrafish mutants. Elife. 11.

Wu T, Hu E, Xu S, Chen M, Guo P, Dai Z, Feng T, Zhou L, Tang W, Zhan L et al. 2021. Clusterprofiler 4.0: A universal enrichment tool for interpreting omics data. Innovation (Camb). 2(3):100141.

Ying J, Srivastava G, Hsieh WS, Gao Z, Murray P, Liao SK, Ambinder R, Tao Q. 2005. The stress-responsive gene gadd45g is a functional tumor suppressor, with its response to environmental stresses frequently disrupted epigenetically in multiple tumors. Clin Cancer Res. 11(18):6442–6449.

Zhang L, Yang Z, Ma A, Qu Y, Xia S, Xu D, Ge C, Qiu B, Xia Q, Li J et al. 2014. Growth arrest and DNA damage 45g down-regulation contributes to janus kinase/signal transducer and activator of transcription 3 activation and cellular senescence evasion in hepatocellular carcinoma. Hepatology. 59(1):178–189.

Zhang P, You N, Ding Y, Zhu W, Wang N, Xie Y, Huang W, Ren Q, Qin T, Fu R et al. 2024. Gadd45g insufficiency drives the pathogenesis of myeloproliferative neoplasms. Nat Commun. 15(1):2989.

