## Supplementary methods and figures for "Genetic context of transgene insertion can influence neurodevelopment in zebrafish"

**Neurobehavioral phenotypes in the pan-neuronal GAL4 zebrafish line *Tg(elavl3:KalTA4)***

**Supplementary Materials and Methods**

**Figure S1:** Additional brain imaging and behavioral phenotypes in *Tg(elavl3:KalTA4)* larvae.

**Figure S2:** Manhattan plots of GO analysis results.

**Figure S3:** Quality control and clustering of sci-RNA-seq data.

**Figure S4:** UMAP representations of expression of select marker genes used to assign neuronal/glial cluster identities.

**Figure S5:** Expression of neuronal genes in the Maturing Zebrafish Telencephalon Atlas.

**Figure S6:** Live confocal images of 6 dpf brains from independently generated *Tol2* and phiC31 *KalTA4* lines crossed with *UAS:GFP* zebrafish.

**Figure S7:** Live fluorescent images of 6 dpf larvae from independently generated *Tol2* and phiC31 *KalTA4* lines crossed with *UAS:GFP* zebrafish.

**Figure S8:** Brain activity and morphology analysis of *pIGLET14a* line.

**Figure S9:** Newly generated Gal4 lines do not exhibit major baseline behavioral phenotypes.

**Supplementary Table 1:** Bulk RNA-seq DESeq2 output of differentially expressed genes.

**Supplementary Table 2:** Gene ontology analysis of upregulated genes.

**Supplementary Table 3:** Gene ontology analysis of downregulated genes.

**Supplementary Table 4:** Gene set enrichment analysis using C5 Molecular Signatures Database.

**Supplementary Table 5:** Gene set enrichment analysis using CNS terms derived from single cell data.

**Supplementary Table 6:** Marker genes used to identify clusters in sci-RNA-seq.

**Supplementary Table 7:** Marker genes used to identify neuronal subclusters in sci-RNA-seq.

**Supplementary Materials and Methods**

Zebrafish husbandry

Both larvae and adult animals were maintained on a 14 hour/10 hour light/dark cycle at 28°C. *Tg(elavl3:KalTA4)* zebrafish (zf562Tg) were maintained as heterozygotes by outcrossing to an Ekkwill (EK)-based wildtype strain. The homozygous *pIGLET14a* line (co2001Tg) was outcrossed to produce heterozygotes, then either incrossed, outcrossed to wild type, or backcrossed to homozygotes for imaging experiments. *Tg(5xUAS:EGFP)* zebrafish (*nkuasgfp1aTg*) were used to visualize *KalTA4* expression in newly generated lines. All experiments were performed at 6 dpf unless otherwise specified. Experimental larvae were raised in 150 mm Petri dishes in fish water with methylene blue at densities of <150 larvae per dish, and debris was removed at 3 dpf. Zebrafish lacking inflated swim bladders were excluded from behavior and imaging experiments with the exception of pERK/tERK immunostaining of *Tg(elavl3:KalTA4)* incrosses. Experiments were conducted blind to genotype, and control larvae were siblings from the same clutch.

Live imaging

Six dpf larval zebrafish were anesthetized with tricaine methanesulfonate (MS-222) and mounted in 2% low melting agarose (Fisher Bioreagents, BP165-25) made in fish water with methylene blue. Side-mounted brightfield images were taken with a Samsung Galaxy S21 and fluorescent images were taken with a Leica M165 FC stereo microscope and K3C color camera. Magnification (3.2x) and exposure (250 ms) were consistent across transgenic lines. Live confocal images were collected using a Zeiss LSM 900 upright confocal microscope with a 20x/1.0 NA water-dipping objective. Zebrafish were pigmented and not treated with PTU.

Brain activity and morphology

pERK/tERK immunostaining and analysis was performed as previously described (Capps et al. 2024). Briefly, 6 dpf larvae were fixed using paraformaldehyde (Polysciences) diluted to 4% in PBS. tERK antibody (Cell Signaling, #4696) was used at a 3:1000 dilution and pERK antibody (Cell Signaling, #4370) was used at a 1:500 dilution. Primary antibody incubation was typically for two days. Images were collected using a Zeiss LSM 900 upright confocal microscope with a 20x/1.0 NA water-dipping objective. Larvae were genotyped after imaging, and confocal stacks were registered to a reference brain using Computational Morphometry Toolkit (CMTK).

Bulk RNA-seq

Bulk RNA-seq was performed as previously described (Capps et al. 2024). Three to four 6 dpf heads were pooled per biological replicate, and four biological replicates were collected per genotype. RNA extraction was performed with E.Z.N.A.® MicroElute® Total RNA Kit (Omega Bio-Tek, R6834-02), and reverse transcription was conducted using Maxima H Minus Reverse Transcriptase (Thermo Scientific, EP0751). Whole-transcriptome amplification was performed with KAPA HiFi HotStart ReadyMix (KAPA Biosystems, KK2601), then PCR products were purified with 0.8X AMPure XP SPRI reagent (Beckman-Coulter, A63881). Library preparation was performed using the Nextera XT DNA Library Preparation Kit (Illumina, FC-131-1096), and libraries were normalized by the UAB Heflin Center for Genomic Science Core Laboratories using qPCR. Sequencing using a NovaSeq 6000 resulted in a typical read depth of 55 million reads per sample.

Bulk RNA-seq analysis

Bulk RNA-seq analysis was performed as previously described (Capps et al. 2024). Reads were aligned using the STAR aligner (2.7.3a-GCC 6.4.0-2.28) (Dobin et al. 2013) to GRCz11 release 104 using the Zebrafish Transcriptome Annotation version 4.3.2 (Lawson et al. 2020). Transcripts with zero counts in four or more samples were removed, and the remaining raw counts were normalized using rlog counts in DESeq2 (Love et al. 2014). DESeq2 script is available at <https://github.com/thymelab/BulkRNASeq>.

Downstream analysis of bulk RNA-seq data was conducted using R (Version 4.3.1). Gene ontology (GO) analysis was performed separately on upregulated genes and downregulated genes with padj <0.05 using the gost function from gprofiler2 (Version 0.2.2) with the false discovery rate correction method (Kolberg et al. 2020). Gene set enrichment analysis (GSEA) was performed as previously described (Capps et al. 2024) using clusterProfiler (Version 4.10.0) (Wu et al. 2021). For chromosomal location GSEA, the gene set was generated using RefSeq Genes and Gene Predictions from the GRCz11 assembly, and genes were sorted by abs(log2 fold change) prior to GSEA. For all other analyses, genes were sorted by log2 fold change. The C5 Molecular Signatures Database (Liberzon et al. 2015) was downloaded using the msigdbr package (Version 7.5.1). The CNS-specific gene set based on 5 dpf single-cell data (Raj et al. 2020) was generated as previously described (Capps et al. 2024). Network plots were created using the emapplot function from enrichplot (Version 1.22.0) and replotted using ggplot2 (Version 3.5.1). Code to reproduce figures is available from Zenodo.

pHH3 staining and analysis

Immunostaining was performed as above using phospho-histone H3 antibody (Cell Signaling, #3377S) and tERK antibody (Cell Signaling, #4696). Images were registered based on tERK staining then segmented with the Z-Brain whole brain mask (Randlett et al. 2015). pHH3 foci in the brain and spinal cord were quantified using the 3D Objects Counter plugin (Bolte and Cordelieres 2006) in Fiji. The script for counting foci is available from Zenodo.

Sci-RNA-seq

Sci-RNA-seq was conducted following the “Tiny-Sci” protocol (Martin et al. 2023). Briefly, *Tg(elavl3:KalTA4)* zebrafish were crossed with *Tg(4xUAS-hmgn6)* zebrafish to produce wildtype, *elavl3:KalTA4*, *4xUAS-hmgn6*, and *elavl3:KalTA4*; *4xUAS-hmgn6* offspring. At 6 dpf, larvae were anesthetized with MS-222 and heads were removed and frozen on dry ice. The remaining body was saved for genotyping. Three heads per biological replicate and three biological replicates per genotype were pooled in 100 µl lysis buffer B in DNA LowBind tubes (Eppendorf, 002431021). Heads were homogenized with a tissue homogenizer at 1 second intervals for 30-45 seconds until no clumps remained. Following fixation with ice-cold methanol and dithiobis (succinimidyl propionate) (DSP, Thermo Fisher, 22586), nuclei were resuspended in sucrose PBS TritonX MgCl_2_ (SPBSTM) and sonicated for 12 seconds on low at 4°C using a Bioruptor® Pico (Diagenode). Nuclei were resuspended in SPBSTM with dNTPs, and 5 ul of each of the twelve samples was pipetted into each well of one column of a twin.tec LoBind PCR plate (Eppendorf, 0030129512). Reverse transcription, ligation, final distribution, second-strand synthesis, protease digestion, tagmentation, PCR amplification, and purification were performed as described except that tagmentation was 10 minutes instead of 5 minutes (Martin et al. 2023). Approximately 1000 nuclei were added to each well during the final distribution step. Plate 1 primer sequences were used for reverse transcription, ligation, and PCR. The library was sequenced on a NextSeq500 by the UAB Heflin Center for Genomic Science Core Laboratories using a 75 cycle kit and 34 cycles for Read1, 10 cycles for Index, and 48 cycles for Read2, resulting in ~36 gigabases of data.

Sci-RNA-seq analysis

Demultiplexing, trimming, alignment, filtering, and gene count processing were performed on the UMass Chan High Performance Computing cluster using scripts available from <https://github.com/bethmartin/sci-RNA-seq3_pipeline/tree/master>. Reads were aligned to GRCz11 release 104 using the Zebrafish Transcriptome Annotation version 4.3.2 (Lawson et al. 2020). Following gene count processing, median UMI count per cell was 365 with 11965 nuclei across all genotypes. Note that the “Tiny-Sci” protocol was not optimized, and further modifications would likely improve the median UMI count and number of nuclei obtained.

Downstream analyses were performed using the Seurat package (Version 5.1.0) (Hao et al. 2024) with R (Version 4.3.3). After filtering to exclude nuclei with <250 or >2500 unique feature counts and nuclei with >5% mitochondrial counts, 5748 nuclei with a median of 424 unique features were retained. Although nuclei of all genotypes were used for clustering, only wildtype and *Tg(elavl3:KalTA4)* nuclei are plotted in Fig. 2. Data were normalized using Seurat’s NormalizeData function, and variable features were selected by combining Seurat’s FindVariableFeatures function with a previously published UMI-based approach (Pandey et al. 2023; Pandey et al. 2018). Data were scaled using the ScaleData function and linear dimensional reduction was performed using the RunPCA function. Cells were clustered using the FindNeighbors function (including dimensions 1 to 15) and the FindClusters function with a resolution of 0.8. Marker genes for clusters were identified using the FindAllMarkers function and cluster identities were defined using canonical marker genes and expression data from Daniocell (Sur et al. 2023). The proportion of cells in each cluster was calculated as previously described (Pandey et al. 2023). The object was then subset to exclude non-neuronal clusters, and reclustered as above using PCA dimensions 1 to 15 and a resolution of 0.7. Seurat objects and code to reproduce figures are available in Zenodo.

Low-pass whole genome sequencing (WGS) and mapping

To purify genomic DNA, *Tg(elavl3:KalTA4)* tissue was incubated overnight at 50°C in extraction buffer containing 10 mM Tris pH 8.2, 10 mM EDTA, 200 mM NaCl, 0.5% SDS, and 200 µg/ml proteinase K. DNA was ethanol precipitated then further purified using the Monarch® Spin gDNA Extraction Kit (NEB, T3010S) according to the manufacturer’s instructions. DNA was sequenced by Genewiz from Azenta Life Sciences to produce ~15 Gb (~10x) low-pass whole genome sequence using Illumina 2x150bp configuration. The grep command was used to identify reads spanning the known *Tol2* sequence and genomic DNA. The insertion site was verified using Sanger sequencing.

qPCR

RNA extraction, cDNA synthesis, and qPCR were performed as previously described (Capps et al. 2024). Three 6 dpf heads were combined and homogenized for each biological replicate, and three biological replicates were collected for each genotype. RNA extraction was performed with E.Z.N.A.® MicroElute® Total RNA Kit (Omega Bio-Tek, R6834-02), and cDNA synthesis was performed with 200 ng RNA input using iScript™ Reverse Transcription Supermix (Bio-Rad, #1708840). 4 ng cDNA input was used in each qPCR reaction, and qPCR was performed using a Bio-Rad CFX96 Touch Real-Time PCR Detection System on a C1000 Touch Thermal Cycler with iTaq™ Universal SYBR® Green Supermix (Bio-Rad, #1725121). Primers for *gadd45ga* spanned exon-exon junctions (forward: 5′-GCTACTACTGGCGATAGAATGC-3′, reverse: 5′-CGCTGTCTGGGTCAACATTC-3′), and primers for the reference gene *rpl13a* were previously published (Rassier et al. 2020).

Cloning plasmids

A phiC31 *attB* site was added to a *Tol2*-compatible backbone that includes the *myl7*:*GFP* transgenesis marker, then putative promoter sequences and *KalTA4* were added to the plasmid using PCR from genomic DNA and Gibson assembly. Promoter for *rtn1a* was ordered as a gBlock (IDT). Full plasmid sequences were verified using Plasmidsaurus. Annotated plasmid maps including full sequences are available in Zenodo.

| **Gene** | **Cloning primers** |
| --- | --- |
| *atp6v0cb* forward | 5′-**AGGGATAACAGGGTAATGATCTAGG**CACATTCAGATGGTCGGGTCA-3′ |
| *atp6v0cb* reverse | 5′-**GCTCGATGGATGAGAGCAGTTTCAT**TATTCCTCTAATTTCACCTCAAAACCC-3′ |
| *elavl3* forward | 5′-**AGGGATAACAGGGTAATGATCTAGG**TTACAAGCTTTATCCAAATTAAGCCA-3′ |
| *elavl3* reverse | 5′-**GCTCGATGGATGAGAGCAGTTTCAT**TCTTGACGTACAAAGATGATAGTGA-3′ |
| *sncb* forward | 5′-**AGGGATAACAGGGTAATGATCTAGG**GTATTGCTTGATCAAAACAACGTCT-3′ |
| *sncb* reverse | 5′-**GCTCGATGGATGAGAGCAGTTTCAT**CTTGGCCCCTCTGCATAAAA-3′ |
| *stmn1b* forward | 5′-**AGGGATAACAGGGTAATGATCTAGG**TTGCCCTCAGTGATAGCAGC-3′ |
| *stmn1b* reverse | 5′-**GCTCGATGGATGAGAGCAGTTTCAT**TGCACTTTCTACTTCGACGGT-3′ |

Generation of transgenic lines

To produce lines using phiC31, *pcDNA3.1 phiC31* plasmid (Bischof et al. 2007; Mosimann et al. 2013) was digested with PvuII-HF, and mRNA was transcribed using the mMESSAGE mMACHINE™ T7 Transcription Kit (Thermo Fisher). mRNA was precipitated and purified with LiCl and ethanol according to the manufacturer’s instructions. Homozygous *pIGLET14a* zebrafish were crossed with *5xUAS:EGFP* zebrafish, and resulting embryos were co-injected with 1 nanoliter of 25 ng/ul phiC31 mRNA and 25 ng/ul plasmid at the one-cell stage. F0 embryos with expression of the *myl7*:*GFP* transgenesis marker were grown to adults and outcrossed to wildtype zebrafish. GFP+ F1 zebrafish were outcrossed to produce larvae for experiments and to confirm integration at the *pIGLET14a* landing site. Integration at the 5′ integration boundary was confirmed using *oRL168 ubi:Switch locus Fw:* 5′CATGATCGAAAACGAGCAATGTC-3′ and *M13 Rev:* 5′-CAGGAAACAGCTATGAC-3′. Integration at the 3′ integration boundary was confirmed using *oRL169 ubi:Switch locus Rev:* 5′GACTGCGTCACTTTGACAACCCTG-3′ and *Tol2 Fw:* 5′AAGTGATCTCCAAAAAATAAGTAC-3′. PCR products for 5′ and 3′ integration boundaries were column purified and sequence was confirmed using Plasmidsaurus.

To produce lines using *Tol2*, embryos were co-injected with 1 nanoliter of 10 ng/ul *Tol2* mRNA (Kawakami et al. 2004) and 20 ng/ul plasmid at the one-cell stage. F0 embryos with expression of the *myl7*:*GFP* transgenesis marker were grown to adults and outcrossed to wildtype zebrafish or *5xUAS:EGFP* zebrafish. GFP+ F1 zebrafish were outcrossed to produce larvae for experiments.

Statistical analysis

Statistical analyses of Mendelian ratio, swim bladder quantification, pHH3 foci, and qPCR were performed with GraphPad Prism (Version 10.4.1). Details about statistical tests and n are available in the figure legends. For brain activity and structure measurements using MapMAPPING (Randlett et al. 2015; Thyme et al. 2019), the significance threshold was based on a false discovery rate where 0.05% of control pixels would be called as significant, and n is available in the figure legends. Statistical analyses for behavioral assays were performed as previously described, and details about the analysis pipeline are available in Joo et al. and Thyme et al (2020; 2019). For both pERK/tERK and behavioral assays, phenotypes were considered significant if they were both statistically significant and repeated in a second independent clutch of larvae. For bulk RNA-seq, the default DESeq2 method was used to determine significance, and both raw and Benjami-Hochberg adjusted P values are available in Supplementary Table 1. For sci-RNA-seq, pairwise two-tailed t-tests were used to detect differences in proportions of cell types using R (Version 4.3.3).

Genotyping

Zebrafish with one copy of *KalTA4* (heterozygous outcross) were genotyped with primers specific to *KalTA4*.

| **Forward** | **Reverse** | **Product Size (bp)** |
| --- | --- | --- |
| 5′-GCTAGAGAGACTGGAACAACTC-3′ | 5′-CGAAACAGTCAGCTGTCTCTGTC-3′ | 262 for *KalTA4* positive |

Zebrafish with one or two copies of *KalTA4* (heterozygous incross) were genotyped with primers specific to *KalTA4* (*KalTA4* product) and to the genomic insertion site (wildtype product).

| **Forward** | **Reverse** | **Product Size (bp)** |
| --- | --- | --- |
| 5′-GCTAGAGAGACTGGAACAACTC-3′ | 5′-CGAAACAGTCAGCTGTCTCTGTC-3′ | 262 for *KalTA4* positive |
| 5′-CCGCTGTCATTTGCAATCTCC-3′ | 5′-GCAATACATTTGTCTCAGGTGTG-3′ | 393 for wild type |

Newly generated *KalTA4* lines were genotyped using three primers per reaction: a forward primer at the 3′ end of the promoter sequence, a reverse primer in *KalTA4* (5′-CAGATGAGCCCTTGTGAGTG-3′, *KalTA4* product), and a reverse primer in the coding sequence of the genomic DNA (wildtype product).

| **Gene** | **Forward (gene-specific)** | **Reverse (gene-specific)** | **Product Size** |
| --- | --- | --- | --- |
| *atp6v0cb* | 5′-TATACGCCTTCGCTCGTGTC-3′ | 5′-CTGAAGACCATCGCTGCTG-3′ | 233 for transgenic; 142 for genomic DNA |
| *elavl3* | 5′-CTGAAGACCTGCAAATCGAAGG-3′ | 5′-CGATTAGACGTGCCAGTATGC-3′ | 216 for transgenic; 259 for genomic DNA |
| *rtn1a* | 5′-ACGACAGCAGTGAGAGAAGC-3′ | 5′-CCGGAGCCTAATCTATCGAATCC-3′ | 189 for transgenic; 328 for genomic DNA |
| *sncb* | 5′-GCTCTCAATTCCAAATTAACCCAC-3′ | 5′-TCGGGCTCCAAGATTCACAG-3′ | 233 for transgenic; 296 for genomic DNA |
| *stmn1b* | 5′-GTGTCTGAAACCGTCGAAGTAG-3′ | 5′-CTTGTCCAGTTCCTTGACCTG-3′ | 192 for transgenic; 210 for genomic DNA |

Zebrafish with one or two copies of the *pIGLET14a* landing site were genotyped with primers specific to *pIGLET14a* (*pIGLET14a* product) and to the genomic insertion site (wildtype product).

| **Forward** | **Reverse** | **Product Size** |
| --- | --- | --- |
| 5′-GCTGGTGCGAATTATTCTGTAGG-3′ | 5′-CCAGTACACGCTACTCAAA-3′ (oRL096 Tol2 remnant Rev) | 631 for presence of *pIGLET14a* landing site |
| 5′-GCTGGTGCGAATTATTCTGTAGG-3′ | 5′-CATGCAAACTCCACACAGAAACG-3′ | 267 for wild type |

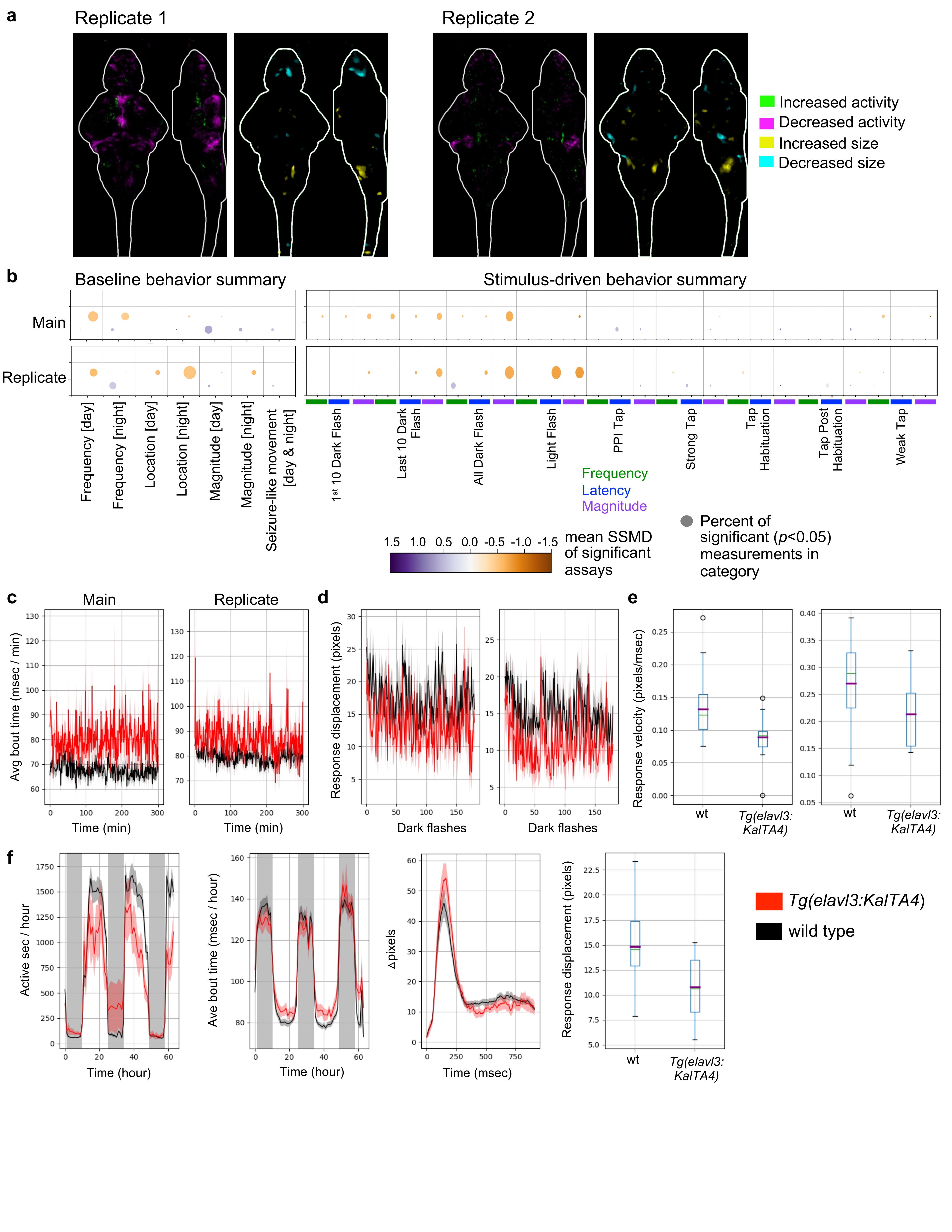

**Fig. S1:** Additional brain imaging and behavioral phenotypes in *Tg(elavl3:KalTA4)* larvae. a) Brain activity and structure maps comparing *Tg(elavl3:KalTA4)* transgenic larvae to wildtype siblings for two additional clutches. Images show sum-of-slices intensity projection inside the brain (white outline). n=19 transgenic and 15 wild type for replicate 1 and 9 transgenic and 20 wild type for replicate 2. b) Summary visualization of the behavioral phenotypes from two independent runs comparing *Tg(elavl3:KalTA4)* transgenic larvae to wildtype siblings. The size of the bubble represents the percent of significant measurements in the summarized category, and the color represents the mean of the strictly standardized mean difference (SSMD) of the significant assays in that category. Detailed behavioral protocols and analysis of the results are available in (Joo et al. 2020; Thyme et al. 2019) with code in <https://github.com/thymelab/ZebrafishBehavior> and <https://github.com/thymelab/DownstreamAnalysis>. n= 17 transgenic and 31 wildtype siblings (main), 14 transgenic and 38 wildtype siblings (replicate). c) Increased bout time for an individual time window (the daytime section at 6 dpf, during the dark flashes or “dfall” window) rather than the entire timecourse shown in the main text and panel S1f. Kruskal-Wallis ANOVA *p* value for main = 0.00011, replicate = 0.029. d) Response displacement for each dark flash event occurring at 6 dpf, shown as a ribbon graph over time. Kruskal-Wallis ANOVA *p* value for main = 0.00014, replicate = 0.00094. e) Response velocity for each all dark flashes occurring at 6 dpf. Kruskal-Wallis ANOVA *p* value for main = 0.0005, replicate = 0.012. f) Replicate graphs for measures shown in Fig. 1. From left to right: 1) Frequency of motion plot, active seconds binned per hour over the entire experiment (Kruskal-Wallis ANOVA *p* value = 0.045). 2) The average time of individual bout motions, binned per hour over the entire experiment (Kruskal-Wallis ANOVA *p* value = 0.63, linear mixed model *p* value = 0.024). This is non-significant with ANOVA because of the timecourse with a very big range of values between day and night. See Thyme et al., 2019 for a description of the linear mixed model and why it is preferable for analyzing significance of the full experimental timecourse. 3) Average movement (pixel-based) of larvae in the mutant and control groups for events where a response to a dark flash was observed. 4) The response displacement for a dark flash response averaged for each zebrafish from all dark flashes (Kruskal-Wallis ANOVA *p* value = 0.00094)

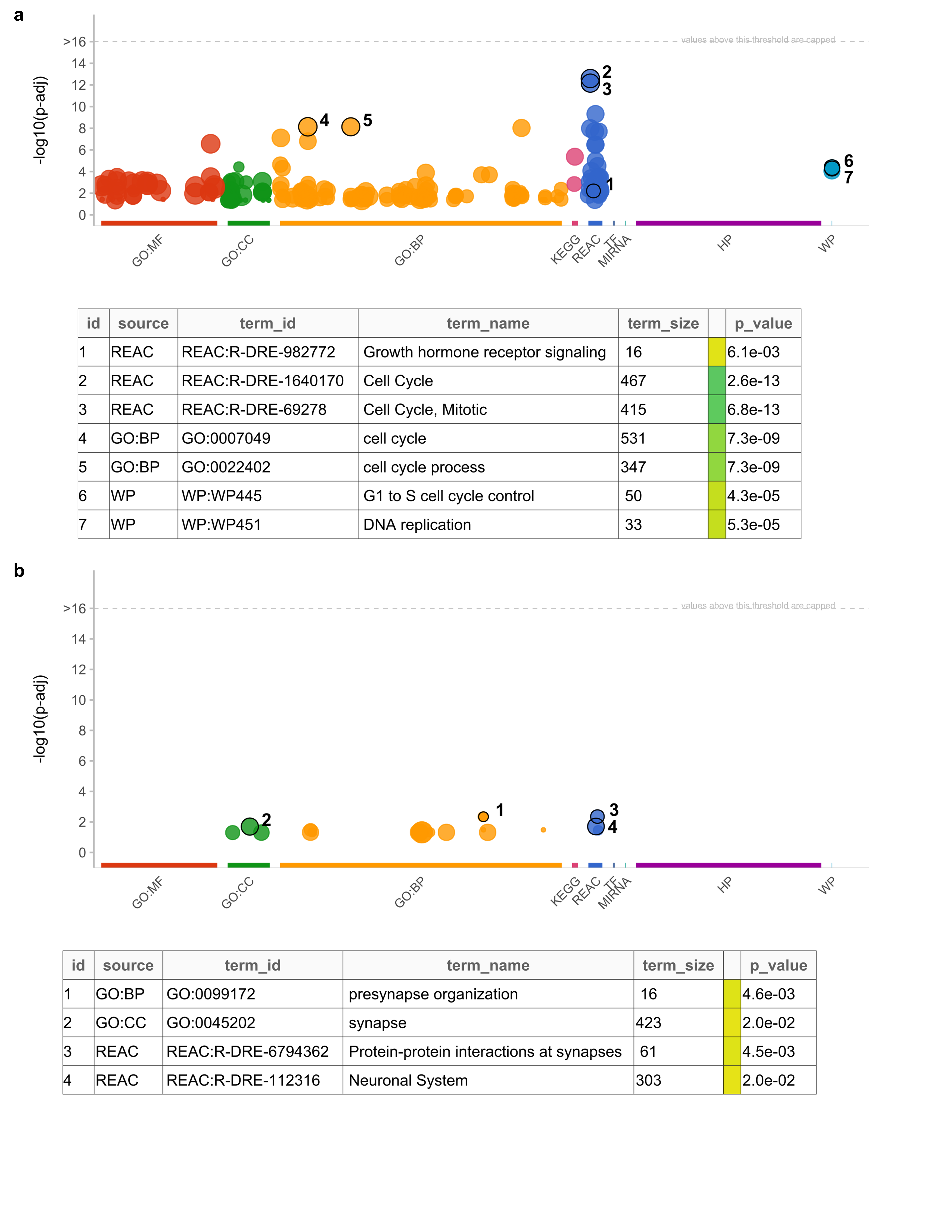

**Figure S2:** Manhattan plots of GO analysis results, with terms derived from GO:MF (molecular function), GO:CC (cellular component), GO:BP (biological process), KEGG (Kyoto Encyclopedia of Genes and Genomes), REAC (Reactome), and WP (WikiPathways). Selected terms are labeled in tables, with full GO results available in Supplementary Tables 2 and 3. Point size represents the number of genes contained in the term. a) Gene ontology analysis using 223 upregulated genes. b) Gene ontology analysis using 111 downregulated genes.

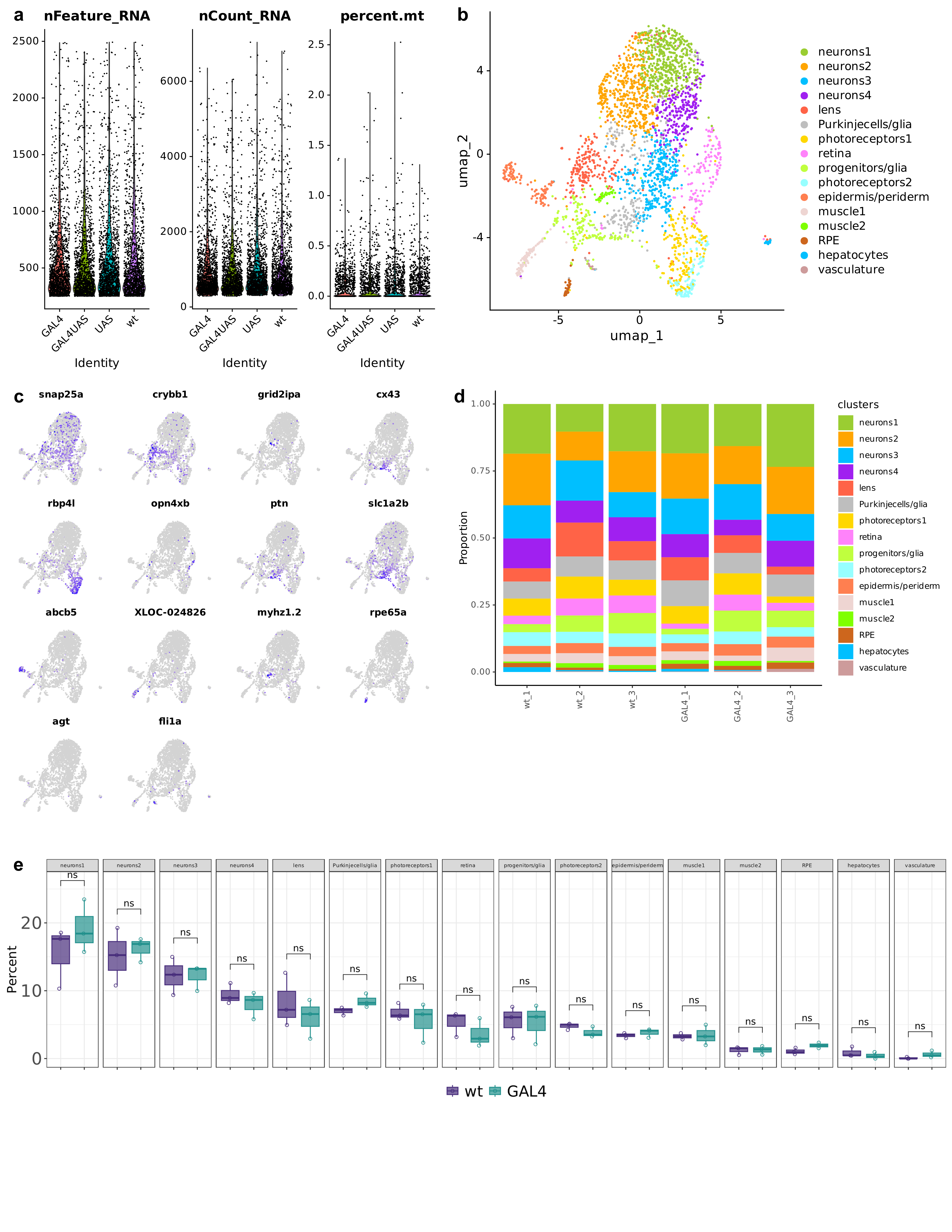

**Figure S3:** Quality control and clustering of sci-RNA-seq data. a) Violin plots showing quality control metrics after filtering for four genotypes across 5748 nuclei. Average unique features per nucleus (nFeature_RNA), molecules per nucleus (nCount_RNA), and percent mitochondrial genes (percent.mt) are 571, 1007, and 0.08, respectively. b) Uniform Manifold Approximation and Projection (UMAP) representation of ten non-neuronal/eye clusters and six neuronal/glial clusters with cells combined from wild type and *Tg(elavl3:KalTA4)* nuclei. c) UMAP representations of expression of select marker genes used to assign cluster identities. Additional marker genes are in Supplementary Table 6. d) Stacked bar plot representing the proportion of cells from each biological replicate contributing to each cluster. n=3 wild type and 3 *Tg(elavl3:KalTa4)*. e) Box plot representing the percentage of cells in each cluster in wild type and *Tg(elavl3:KalTa4)*. *Tg(elavl3:KalTa4)* does not differ significantly from wild type for any cluster. Statistics, two-sided t-test.

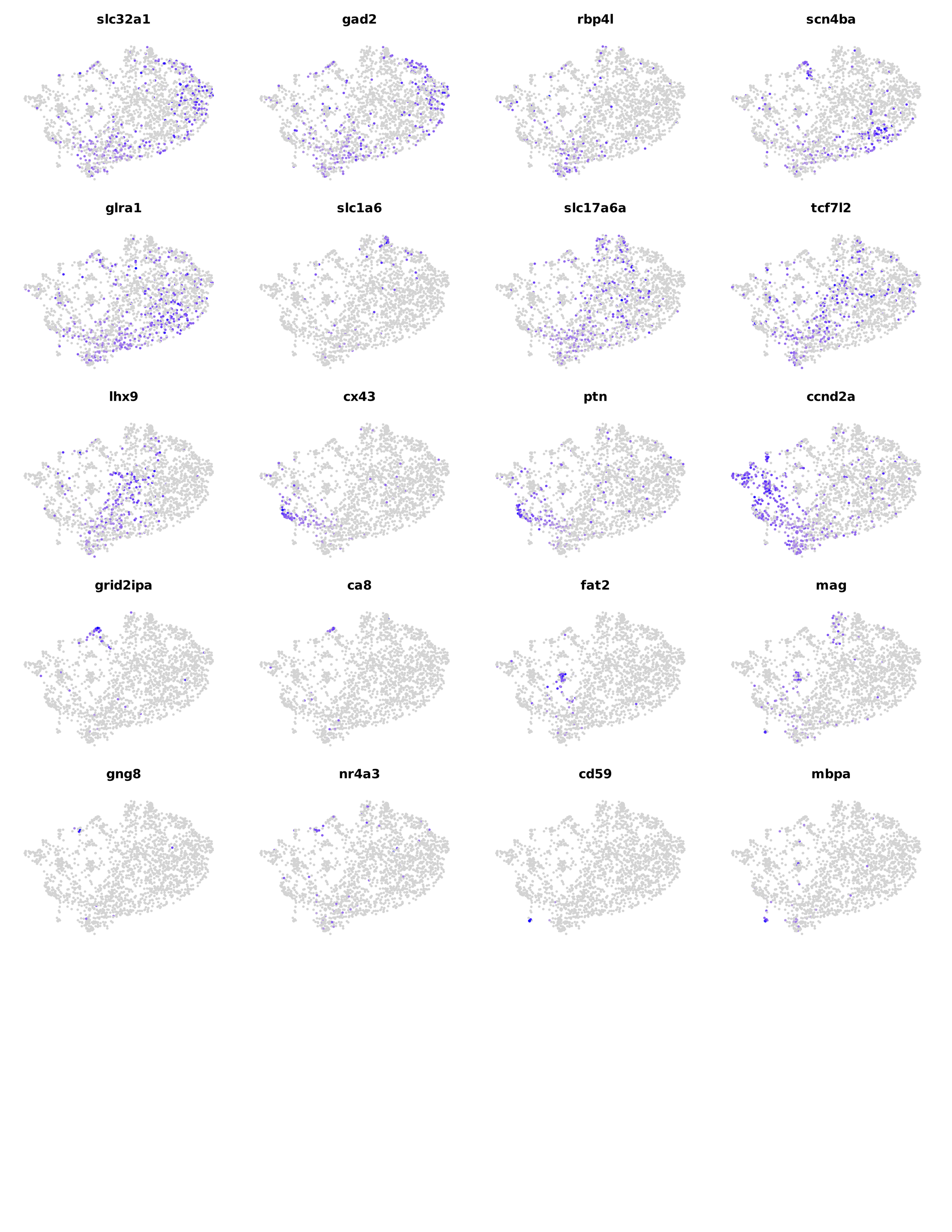

**Figure S4:** UMAP representations of expression of select marker genes used to assign neuronal/glial cluster identities. Additional marker genes are in Supplementary Table 7.

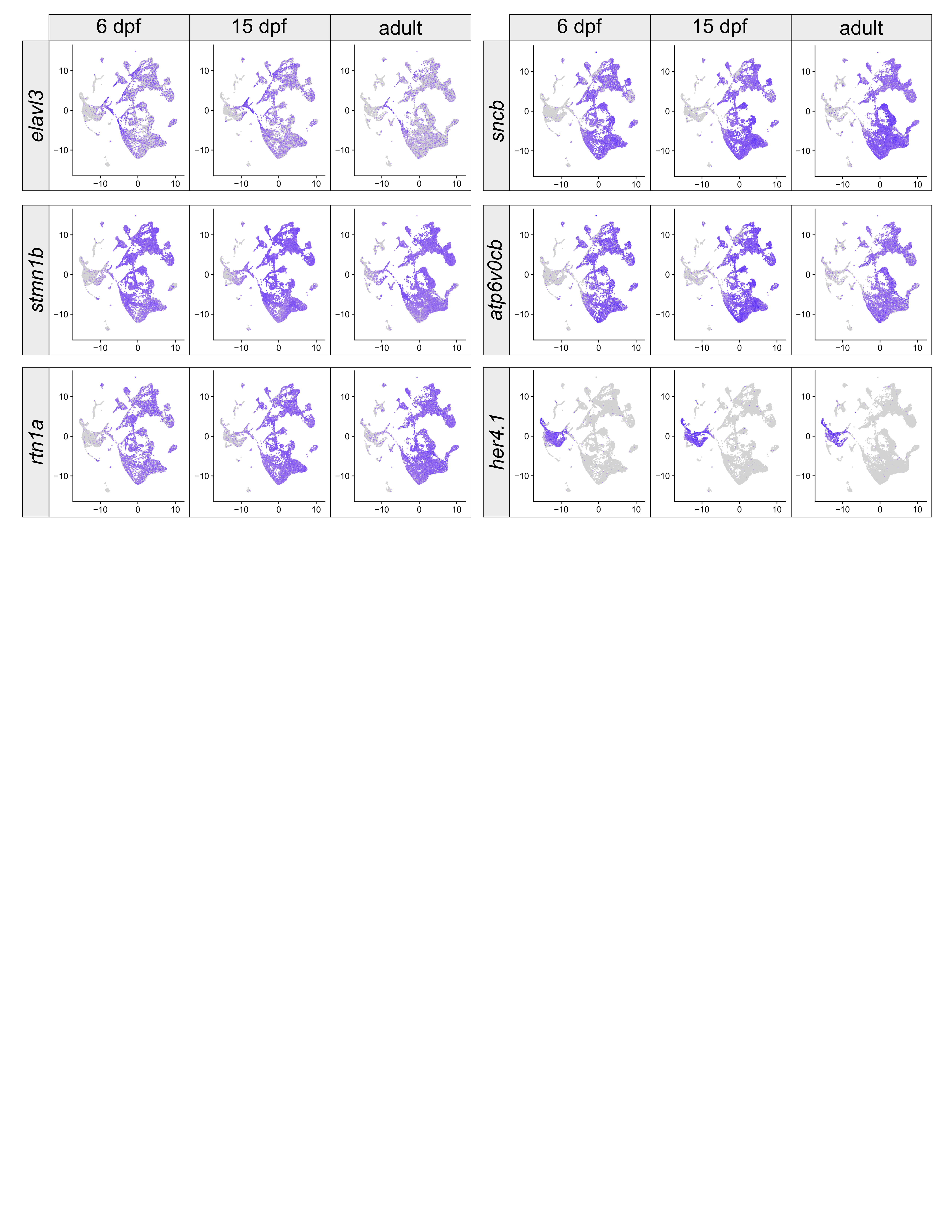

**Figure S5:** Expression of neuronal genes in the Maturing Zebrafish Telencephalon Atlas obtained from <https://zfforebrain.thymelab.org/> (Pandey et al. 2023). UMAP representations of gene expression, with gray indicating undetectable expression and purple indicating high expression. *Elavl3* is expressed in neuronal precursors and mature neurons with peak expression in committed precursors and relatively weaker expression in mature adult neurons. *Stmn1b* and *rtn1a* show similar expression patterns with some expression in progenitors and committed precursors but with high expression maintained in mature adult neurons. *Sncb* and *atp6v0cb* have relatively low expression in progenitors with high expression in mature neurons that is maintained into adulthood. For comparison, expression of *her4.1* is high in progenitors but turns off as progenitors commit to neuronal differentiation.

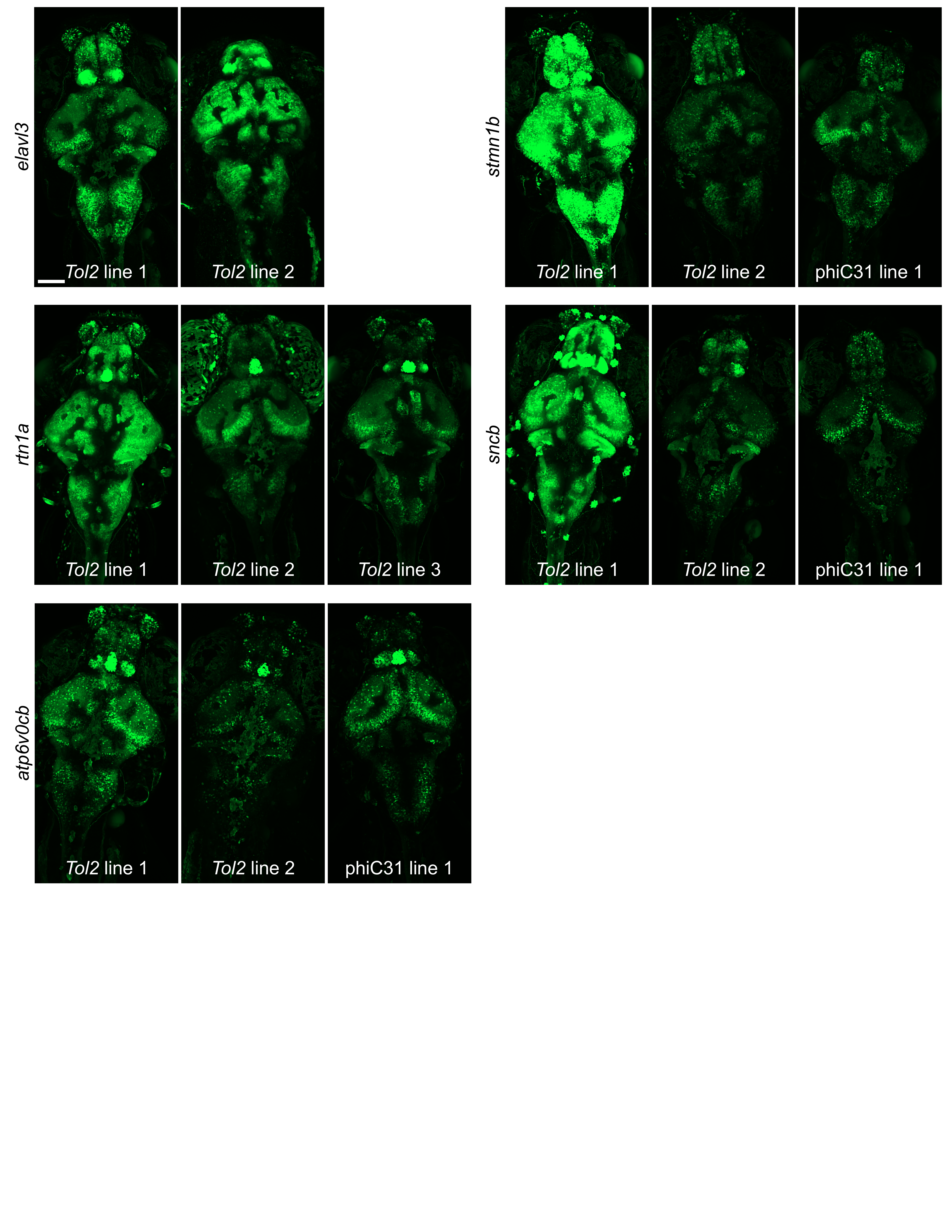

**Figure S6:** Live confocal images of 6 dpf brains from independently generated *Tol2* and phiC31 *KalTA4* lines crossed with *UAS:GFP* zebrafish. Lines show variation in fluorescence intensity both within and between promoters, with *atp6v0cb* lines showing relatively low expression of GFP. Images shown are 145-slice maximal projections with 3 μm interval. Scale bar = 100 μm.

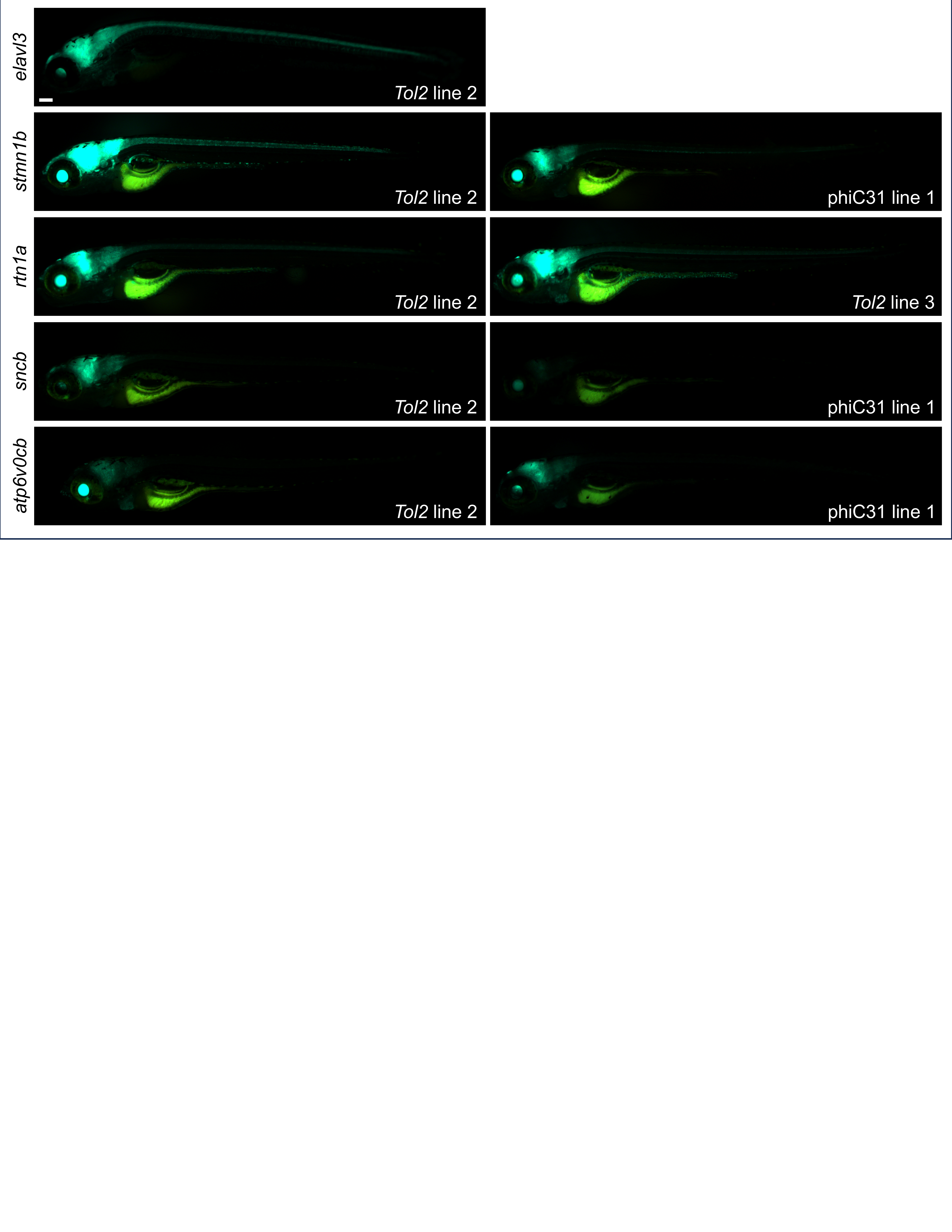

**Figure S7:** Live fluorescent images of 6 dpf larvae from independently generated *Tol2* and phiC31 *KalTA4* lines crossed with *UAS:GFP* zebrafish. Autofluorescence appears in the yolk. Scale bar = 100 μm.

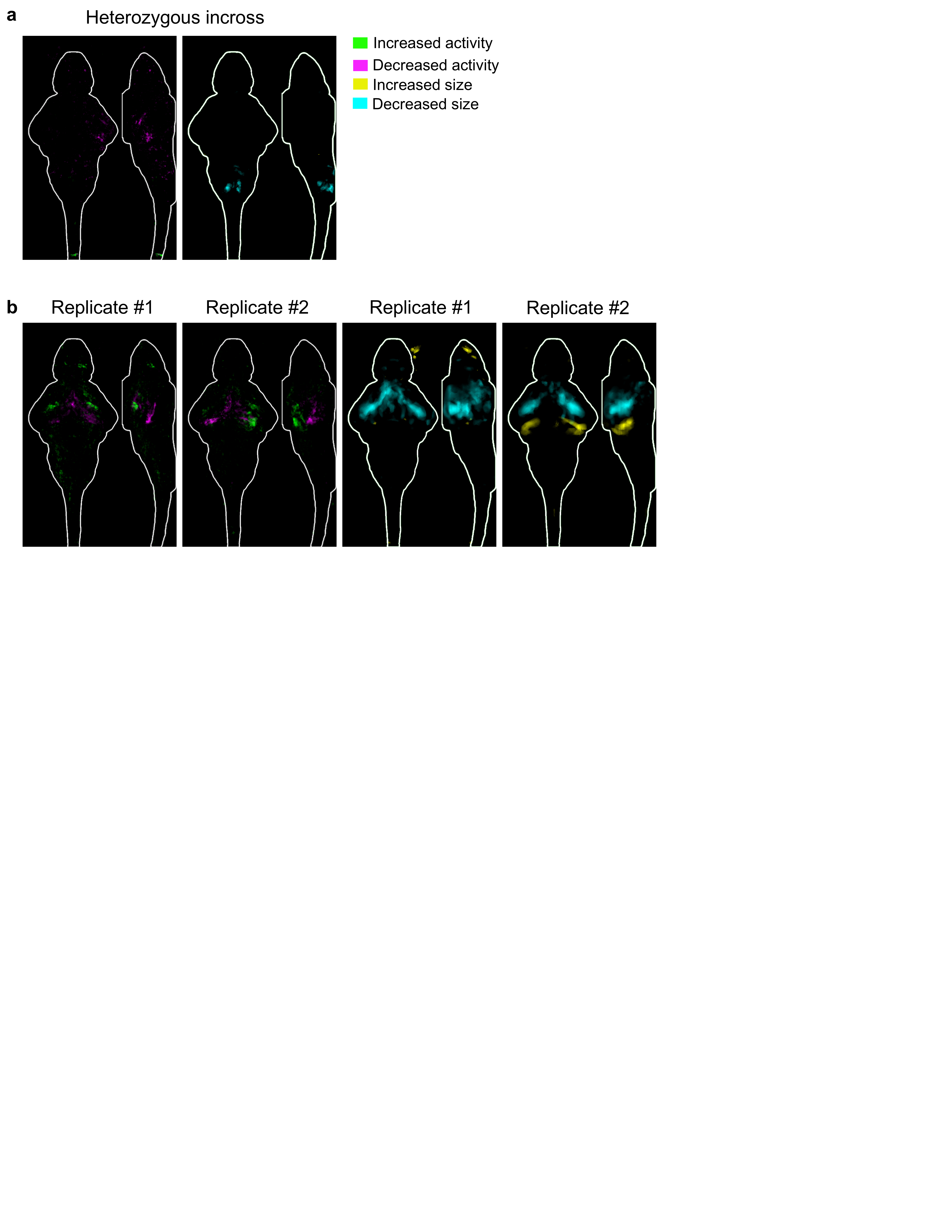

**Figure S8:** Brain activity and morphology analysis of *pIGLET14a* line. Images show sum-of-slices intensity projection inside the brain (white outline). a) Brain activity and structure maps comparing heterozygous *pIGLET14a* transgenic larvae to wildtype siblings from a heterozygous incross (n=13 heterozygous and 11 wild type). Maps show minimal phenotypes in heterozygous zebrafish. b) Brain activity and structure maps comparing homozygous *pIGLET14a* transgenic larvae to heterozygous siblings from homozygous x heterozygous backcross. Two independent replicates show decreased size of the hypothalamus in homozygous zebrafish. This phenotype was not replicated in homozygous fish derived from a heterozygous incross, suggesting that it may derive from incrossing/backcrossing. n= 24 homozygous and 36 heterozygous for replicate 1 and 23 homozygous and 36 heterozygous for replicate 2.

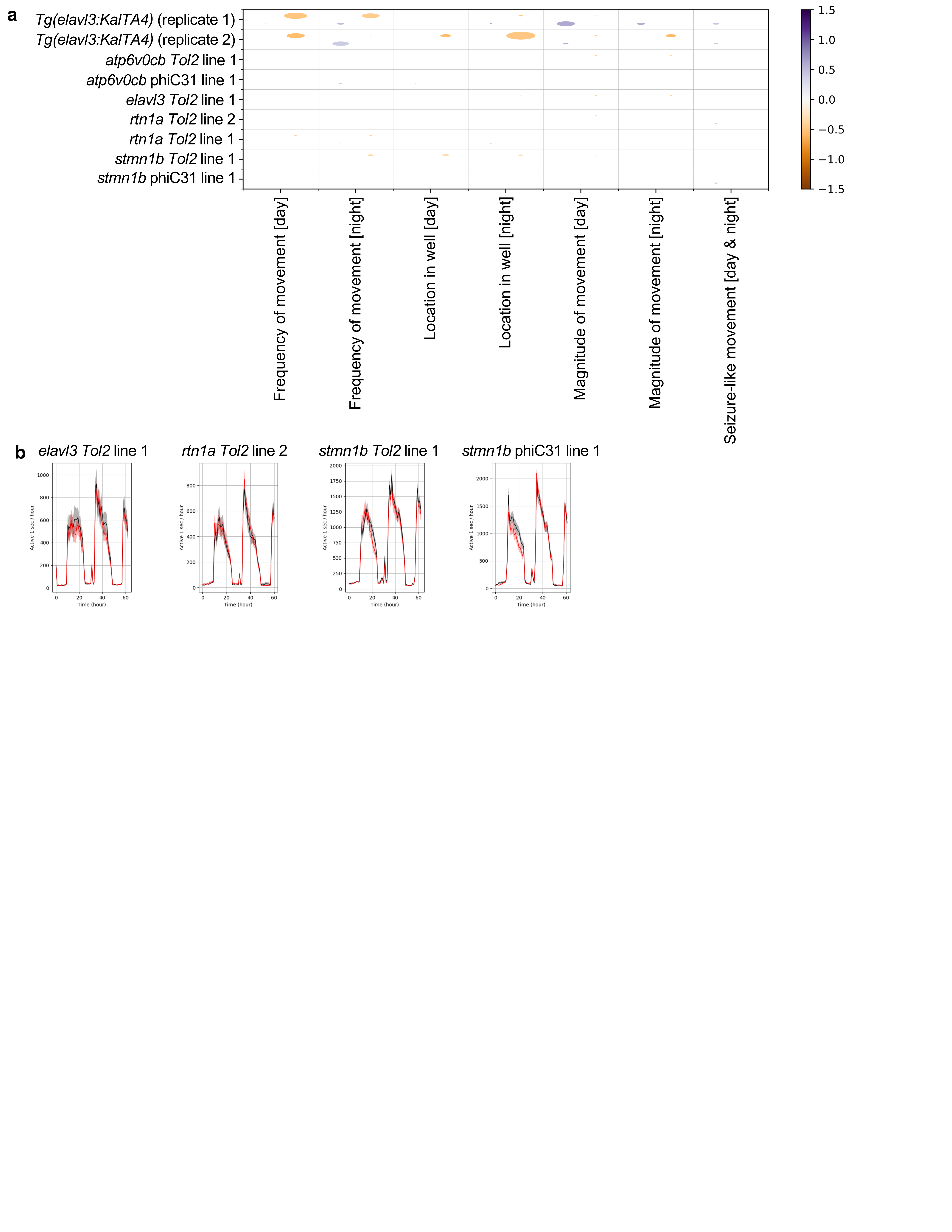

**Figure S9:** Newly generated Gal4 lines do not exhibit major baseline behavioral phenotypes. a) Summary visualization of the baseline behavioral phenotypes from two independent runs comparing original *Tg(elavl3:KalTA4)* transgenic larvae to wildtype siblings (top two rows, data also in Supplementary Fig. S1) and new transgenic lines to wildtype siblings. The size of the bubble represents the percent of significant measurements in the summarized category, and the color represents the mean of the strictly standardized mean difference (SSMD) of the significant assays in that category. n= 17 transgenic and 31 wildtype siblings (*Tg(elavl3:KalTA4)* replicate 1), 14 transgenic and 38 wildtype siblings (*Tg(elavl3:KalTA4)* replicate 2), 56 transgenic and 35 wildtype siblings (*atp6v0cb Tol2* line 1), 24 transgenic and 29 wildtype siblings (*atp6v0cb* phiC31 line 1), 37 transgenic and 30 wildtype siblings (*elavl3 Tol2* line 1), 29 transgenic and 44 wildtype siblings (*rtn1a Tol2* line 2), 43 transgenic and 19 wildtype siblings (*rtn1a Tol2* line 1), 41 transgenic and 55 wildtype siblings (*stmn1b Tol2* line 1), and 37 transgenic and 47 wildtype siblings (*stmn1b* phiC31 line 1). b) Representative frequency of motion plots, with active seconds binned per hour over the entire experiment. Red line represents transgenic larvae and black line represents wildtype controls.
